# Constraints on the evolution of toxin-resistant Na,K-ATPases have limited dependence on sequence divergence

**DOI:** 10.1101/2021.11.29.470343

**Authors:** Shabnam Mohammadi, Santiago Herrera-Álvarez, Lu Yang, María del Pilar Rodríguez-Ordoñez, Karen Zhang, Jay F. Storz, Susanne Dobler, Andrew J. Crawford, Peter Andolfatto

**Affiliations:** School of Biological Sciences, University of Nebraska, Lincoln, NE, USA; Molecular Evolutionary Biology, Institute of Zoology, Universität Hamburg, Hamburg, Germany; Department of Ecology and Evolution, Princeton University, Princeton, NJ, USA; Department of Biological Sciences, Universidad de los Andes, Bogotá, 111711, Colombia; Department of Ecology and Evolution, University of Chicago, Chicago, IL, USA; Department of Biological Sciences, Columbia University, New York, NY, USA

**Author notes:** Co-first authorship (Shabnam Mohammadi, Santiago Herrera-Álvarez, Lu Yang). Université Paris-Saclay Evry, Evry, France (María del Pilar Rodríguez-Ordoñez). **Author Contributions** PA and AJC conceived of and oversaw the project; SM, JFS, SD, AJC and PA designed experiments; KZ, LY, MPRO, SHA, SM collected data; SM, SHA and PA performed evolutionary and statistical analyses; SM, SHA, and PA wrote the paper; All authors edited the manuscript.

**Keywords:** Epistasis, protein evolution, cardiotonic steroids, toxin resistance, adaptation

## Abstract

A growing body of theoretical and experimental evidence suggests that intramolecular epistasis is a major determinant of rates and patterns of protein evolution and imposes a substantial constraint on the evolution of novel protein functions. Here, we examine the role of intramolecular epistasis in the recurrent evolution of resistance to cardiotonic steroids (CTS) across tetrapods, which occurs via specific amino acid substitutions to the α-subunit family of Na,K-ATPases (ATP1A). After identifying a series of recurrent substitutions at two key sites of ATP1A that are predicted to confer CTS resistance in diverse tetrapods, we then performed protein engineering experiments to test the functional consequences of introducing these substitutions onto divergent species backgrounds. In line with previous results, we find that substitutions at these sites can have substantial background-dependent effects on CTS resistance. Globally, however, these substitutions also have pleiotropic effects that are consistent with additive rather than background-dependent effects. Moreover, the magnitude of a substitution’s effect on activity does not depend on the overall extent of ATP1A sequence divergence between species. Our results suggest that epistatic constraints on the evolution of CTS-resistant forms of Na,K-ATPase likely depend on a small number of sites, with little dependence on overall levels of protein divergence. We propose that dependence on a limited number sites may account for the observation of convergent CTS resistance substitutions observed among taxa with highly divergent Na,K-ATPases.

**Significance Statement:** Individual amino acid residues within a protein work in concert to produce a functionally coherent structure that must be maintained even as orthologous proteins in different species diverge over time. Given this dependence, we expect identical mutations to have more similar effects on protein function in more closely related species. We tested this hypothesis by performing protein-engineering experiments on ATP1A, an enzyme mediating target-site insensitivity to cardiotonic steroids (CTS) in diverse animals. These experiments reveal that the phenotypic effects of substitutions can sometimes be background-dependent, but also that the magnitude of these phenotypic effects does not correlate with overall levels of ATP1A sequence divergence. Our results suggest that epistatic constraints are determined by states at a small number of sites, potentially explaining the frequent convergent CTS resistance substitutions among Na,K-ATPases of highly divergent taxa.

## Introduction

Instances of parallel and convergent (hereafter “convergent”) evolution represent a useful paradigm to examine the factors that limit the rate of adaptation and the extent to which adaptive evolutionary paths are predictable [1,2]. The evolution of resistance to cardiotonic steroids (CTS) in animals represents one of the clearest examples of convergent molecular evolution. CTS are potent inhibitors of Na,K-ATPase (NKA, Fig. 1A), a protein that plays a critical role in maintaining membrane potential and is consequently vital for the maintenance of many physiological processes and signaling pathways in animals [3]. CTS inhibit NKA function by binding to a highly conserved domain of the protein’s α-subunit (ATP1A) and blocking the exchange of Na^+^ and K^+^ ions [3]. Thus, NKA is often the target of convergent evolution of CTS resistance across widely divergent species, including insect herbivores that feed on toxic plants [4,5] as well as predators that feed on toxic prey [6–10].

**Figure 1.**
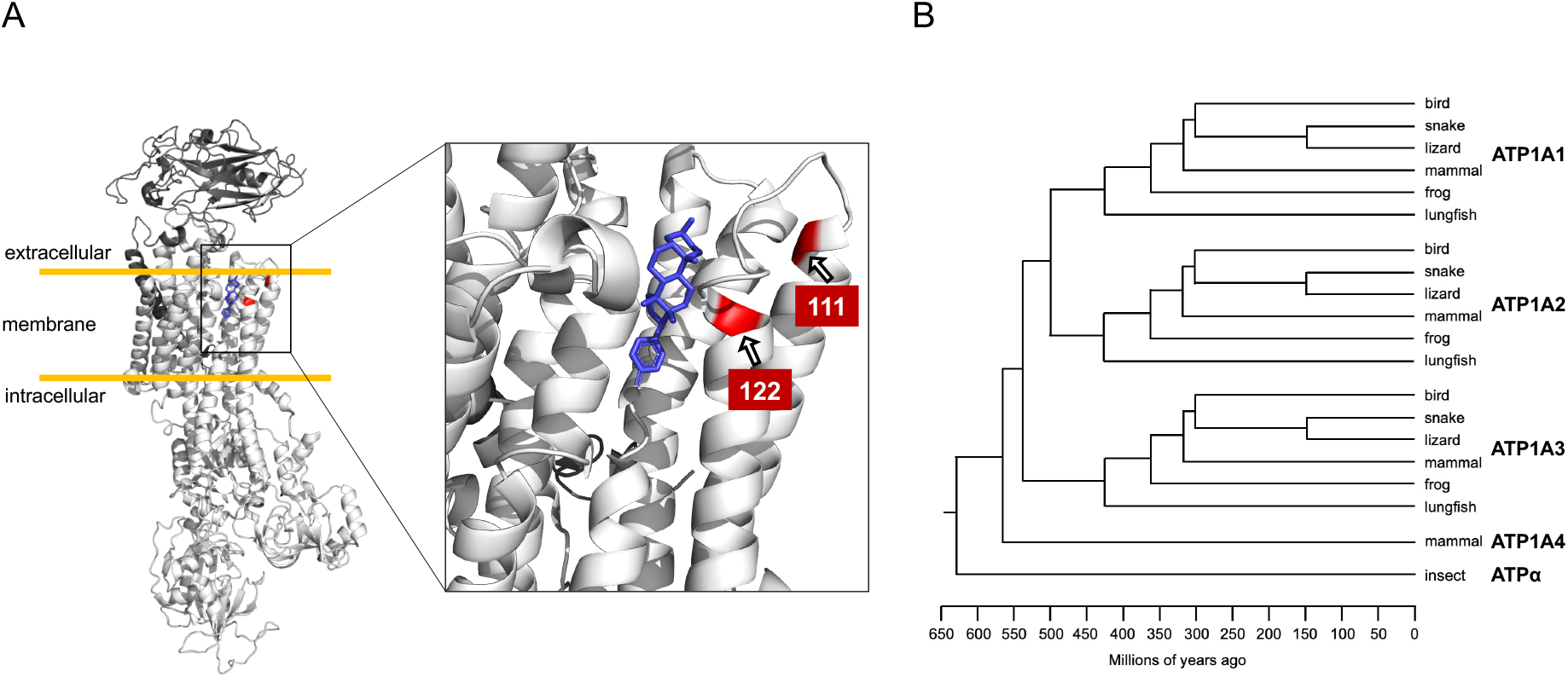
Na,K-ATPase structure and phylogenetic relationships of ATP1A paralogs among vertebrates. (A) Crystal structure of an Na^+^,K^+^-ATPase (NKA) with a bound the representative CTS bufalin in blue (PDB 4RES). The zoomed-in panel shows the H1-H2 extracellular loop, highlighting two amino acid positions (111 and 122 in red) that have been implicated repeatedly in CTS resistance. We highlight key examples of convergence in amino acid substitutions at sites in the H1-H2 extracellular loop associated with CTS resistance in Fig. 3. (B) Phylogenetic relationships among ATP1A paralogs of vertebrates and ATPα of insects.

Patterns of convergence in adaptive protein evolution are influenced by the mutational target size (i.e. the number of potentially adaptive mutations), the degree of pleiotropy (i.e. the effect of a given mutation on multiple phenotypes), and intramolecular epistasis (i.e. nonadditive interactions between mutant sites in the same protein) [11–19]. If the phenotypic and fitness effects of mutations depend on the protein sequence background on which they arise (i.e. there is intramolecular epistasis), a given mutation is expected to have more similar phenotypic and fitness effects in orthologs from closely-related species. Therefore, the probability of convergent substitution is expected to decrease with increasing sequence divergence between orthologous proteins in different species. Consistent with this expectation, such a decline is observed in broad-scale phylogenetic comparisons of mitochondrial [20] and nuclear [21,22] proteins. While these results suggest epistasis is an important global determinant of patterns of convergent protein evolution, studies linking these broad-scale observations with functional data at the level of individual proteins are lacking.

Functional investigations of CTS resistance-conferring substitutions at two key sites (111 and 122) in ATP1A orthologs of Drosophila [23,24] and Neotropical grass frogs [25] revealed associated negative pleiotropic effects on protein function and showed that evolution at other sites in the protein mitigate these detrimental effects. In light of these pleiotropic and epistatic constraints, it is curious that convergent CTS-resistant substitutions are often observed among ATP1A orthologs of highly divergent species. Due to limited comparative functional data, the generality of pleiotropic and epistatic constraints on ATP1A-mediated CTS resistance, and specifically the predicted dependence on evolutionary distance, remain poorly understood.

To achieve a clearer picture of the phylogenetic distribution of CTS resistance substitutions, a more complete and consistent sampling of ATP1A is needed in vertebrates. Broad phylogenetic comparisons in vertebrates have primarily focused on the H1-H2 extracellular loop of ATP1A, which represents only a subset of the CTS-binding domain. Further, most vertebrates possess three paralogs of ATP1A that have different tissue-specific expression profiles and are associated with distinct physiological roles (Fig. 1B) [3,26]. Previous studies of the ATP1A paralogs of vertebrate taxa focused on ATP1A3 in reptiles [7,8,27,28], ATP1A1 and/or ATP1A2 in birds and mammals [10,28], and either ATP1A1 or ATP1A3 in amphibians [6,28]. We therefore lack a comprehensive and systematic survey of amino acid variation in the ATP1A protein family across vertebrates.

To bridge this gap, we first surveyed variation in near full-length coding sequences of the three NKA α-subunit paralogs (ATP1A1, ATP1A2, ATP1A3) that are shared across major extant tetrapod groups (mammals, birds, non-avian reptiles, and amphibians), and identified substitutions that occur repeatedly among divergent lineages. If the phenotypic effects of these substitutions depend on states at a large number of sites throughout the protein, we expect that identical substitutions should have increasingly distinct effects on more highly divergent proteins. Focusing on two key sites implicated in CTS resistance across animals (111 and 122), we tested whether substitutions at these sites have increasingly distinct phenotypic effects on more divergent genetic backgrounds. We engineered several common substitutions at sites 111 and 122 of ATP1A1 that differ between species to reveal potential ‘cryptic’ epistasis [16,29]. By quantifying the level of CTS resistance conferred by these substitutions on different backgrounds, as well as their pleiotropic effects on enzyme function, we evaluate the extent to which overall protein sequence background has constrained the evolution of CTS-resistant forms of ATP1A1 across tetrapods.

## Results

### Patterns of ATP1A sequence evolution across species and paralogs

To obtain a more comprehensive portrait of ATP1A amino acid variation among tetrapods, we created multiple sequence alignments for near full-length ATP1A proteins for the three ATP1A paralogs shared among vertebrates. In addition to publicly available data, we generated new RNA-seq data for 27 non-avian reptiles (PRJNA754197) (Table S1-S2). We then *de novo* assembled full-length transcripts of all ATP1A paralogs using these and generated new RNA-seq data for 18 anuran species [25] (PRJNA627222) to achieve better representation for these groups. In total, this dataset comprises 429 species for ATP1A1, 197 species for ATP1A2 and 204 species for ATP1A3 (831 sequences total, including the newly generated data; Supplemental Dataset 1, Fig. S1).

Our survey reveals numerous substitutions at sites implicated in CTS resistance of NKA (Fig. 2; Supplementary Dataset 2; for comparison to insects, see Supplemental file 1 of ref. [23]). As anticipated from studies of full-length sequences in insects [4,5,23], most amino acid variation among species and paralogs is concentrated in the H1-H2 extracellular loop (residues 111-122; Fig 1A). Despite harboring just 28% of 43 sites previously implicated in CTS resistance [30], the H1-H2 extracellular loop contains 81.4% of all substitutions identified among the three ATP1A paralogs (Fig. S2).

**Figure 2.**
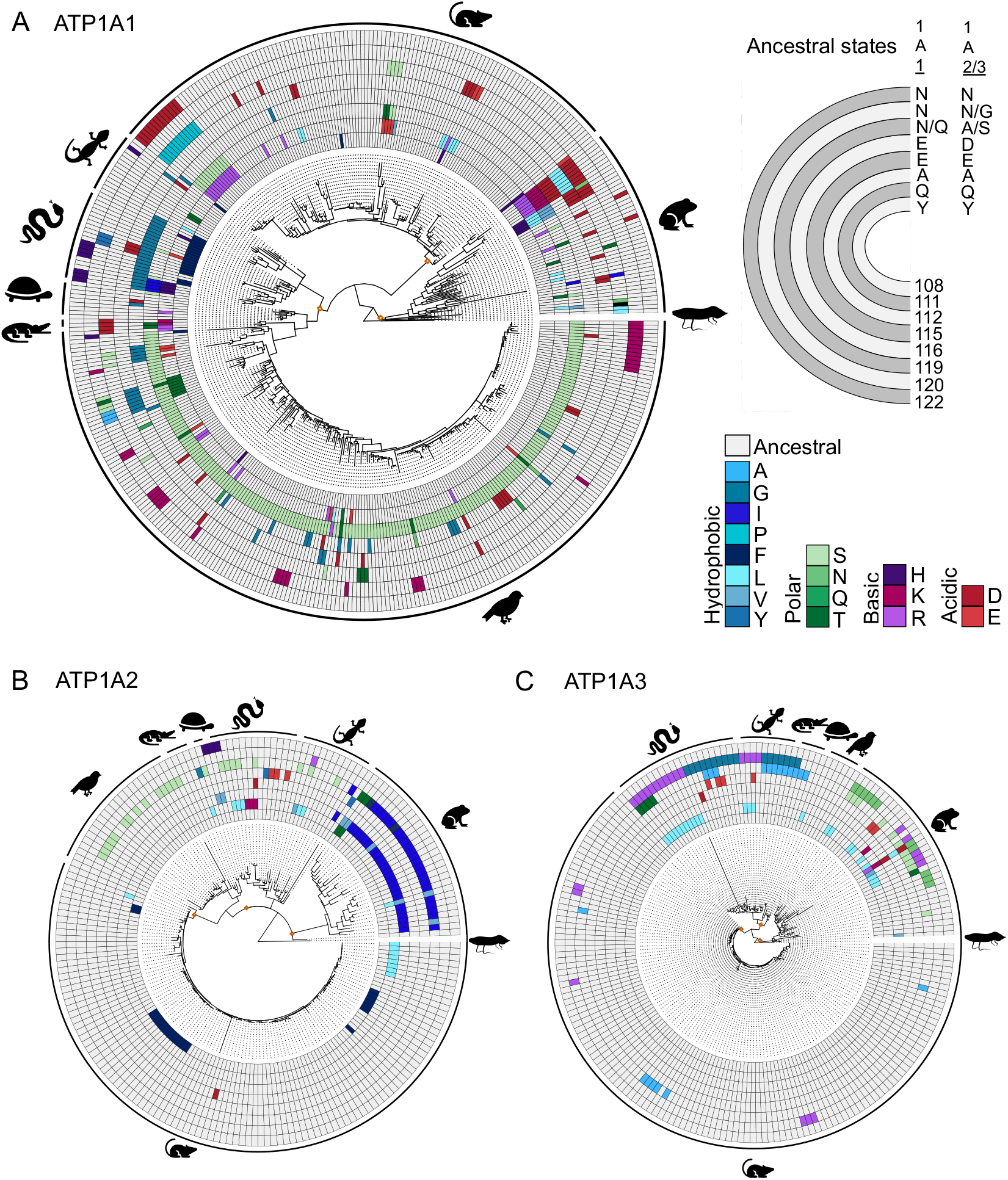
Patterns of molecular evolution in the α(H1–H2) extracellular loop of ATP1A paralogs shared among tetrapods. (A) Maximum likelihood phylogeny of tetrapod ATP1A1, (B) ATP1A2, and (C) ATP1A3. The character states for eight sites relevant to CTS resistance in and near the H1-H2 loop of the NKA protein are shown at the node tips. Yellow internal nodes indicate ancestral sequences reconstructed to infer derived amino acid states across clades to ease visualization; nodes reconstructed: most recent common ancestor (MRCA) of mammals, of reptiles, and of amphibians. *Top right*, each semi-circle indicates the site mapped in the main phylogeny with the inferred ancestral amino acid state for each of the three yellow nodes (posterior probability >0.8). In ATP1A1, site 119 was inferred as Q119 for amphibians and mammals, and N119 for reptiles (Table S6); in ATP1A2-3 site 119 was inferred as A119 for amphibians and reptiles, and S119 for mammals (Table S6). Site number corresponds to pig (*Sus scrofa*) reference sequence. Higher number and variation of substitutions in ATP1A1 stand out in comparison to the other paralogs.

Our survey reveals several clade- and paralog-specific patterns. Notably, ATP1A1 exhibits more variation among species at sites implicated in CTS resistance (Fig. 2). Most of the variation in ATP1A2 at these sites is restricted to squamate reptiles and ATP1A3 lacks substitutions at site 122 altogether, despite the well-known potential for substitutions at this site to confer CTS resistance [25,31]. Looking across species and paralogs, the extent of convergence at sites 111 and 122 is remarkable (Figs. 2-3): for example, the substitutions Q111E, Q111T, Q111H, Q111L, and Q111V all occur in parallel in multiple species of both insects and vertebrates. N122H and N122D also frequently occur in parallel in both of these major clades. The frequent convergence of CTS-sensitive (i.e. Q111 and N122) to CTS-resistant states at these sites has been interpreted as evidence for adaptive significance of these substitutions [4–7], but may also reflect mutational biases [32] and the nature of physico-chemico constraints [21,33].

In contrast, some convergence is restricted to specific clades: for example, Q111R occurs in parallel across tetrapods but has not been observed in insects. Similarly, the combination Q111R+N122D has evolved three times independently in ATP1A1 of tetrapods but is not observed in insects. Conversely, insects have evolved Q111V+N122H independently four times, but this combination has never been observed in tetrapods. These patterns suggest that the fitness effects of some CTS-resistant substitutions depend on genetic background (i.e. epistatic constraints), with the result that CTS-resistance evolved via different mutational pathways in different lineages.

Beyond known CTS-resistant substitutions at sites 111 and 122, some taxa have evolved other paths to CTS resistance. For example, horned frogs of the genus *Ceratophyrs* are known to prey on CTS-containing toads [34] and their ATP1A1 harbors a known CTS-resistant substitution at site 121 (D121N, Supplementary Dataset 2). This substitution is rare among vertebrates but has been previously reported in CTS-adapted milkweed bugs [4,5]. Similarly, the known CTS resistance substitution C104Y is observed in ATP1A1 among garter snakes of the genus *Thamnophis* (Supplementary Dataset 2) and CTS-adapted milkweed weevils [5]. Histricognathi rodents, including Chinchilla (*Chinchilla lanigera*), and yellow-throated sandgrouse (*Pterocles gutturalis*) show distinct single-amino acid insertions in the H1–H2 extracellular loop, a characteristic that has been previously associated with CTS resistance in pyrgomorphid grasshoppers [30,35]. Further, *in lieu* of variation at site 122, ATP1A3 of tetrapods harbors frequent convergent substitutions at site 120 (G120R, see also [7]). Interestingly, this site also shows substantial convergent substitution in the ATP1A1 paralog of birds (where N120K occurs eight times independently) but is mostly invariant in ATP1A1 of other tetrapods.

### Context-dependent CTS resistance of substitutions at sites 111 and 122

The clade- and paralog-specific patterns of substitution among ATP1A paralogs outlined above suggest that the evolution of CTS resistance may be highly dependent on sequence context. However, the functional effects of the vast majority of these substitutions on the diverse genetic backgrounds in which they occur remain largely unknown [25,27,31]. Given the diversity and broad phylogenetic distribution of convergent substitutions at sites 111 and 122, and the documented effects of some of these substitutions on CTS resistance, we experimentally tested the extent to which functional effects of substitutions at these sites are background-dependent.

We focused functional experiments on ATP1A1, because it is the most ubiquitously expressed paralog and exhibits both the most sequence diversity and the broadest phylogenetic distribution of convergent substitutions. Specifically, we considered ATP1A1 orthologs from nine representative tetrapod species that possess different combinations of wild-type amino acids at 111 and 122 (Fig. 4A). Our taxon sampling included two lizards, two snakes, two birds, two mammals, and previously published data for one amphibian (Fig. S6; Fig. S7; Table S3). The ancestral amino acid states of sites 111 and 122 in tetrapods are Q and N, respectively. We found that the sum of the number of derived states at positions 111 and 122 is a strong predictor of the level of CTS-resistance (Fig 4B, IC_50_, Spearman’s *r*_S_=0.85, p=0.001). Nonetheless, we also found greater than 10-fold variation in CTS resistance among enzymes that had identical paired states at 111 and 122 (e.g., compare chinchilla (CHI) versus red-necked keelback snake (KEE) or compare rat (RAT) versus the resistant paralog of grass frog (GRA_R_)). These differences suggest that substitutions at other sites also contribute to CTS resistance.

**Figure 3.**
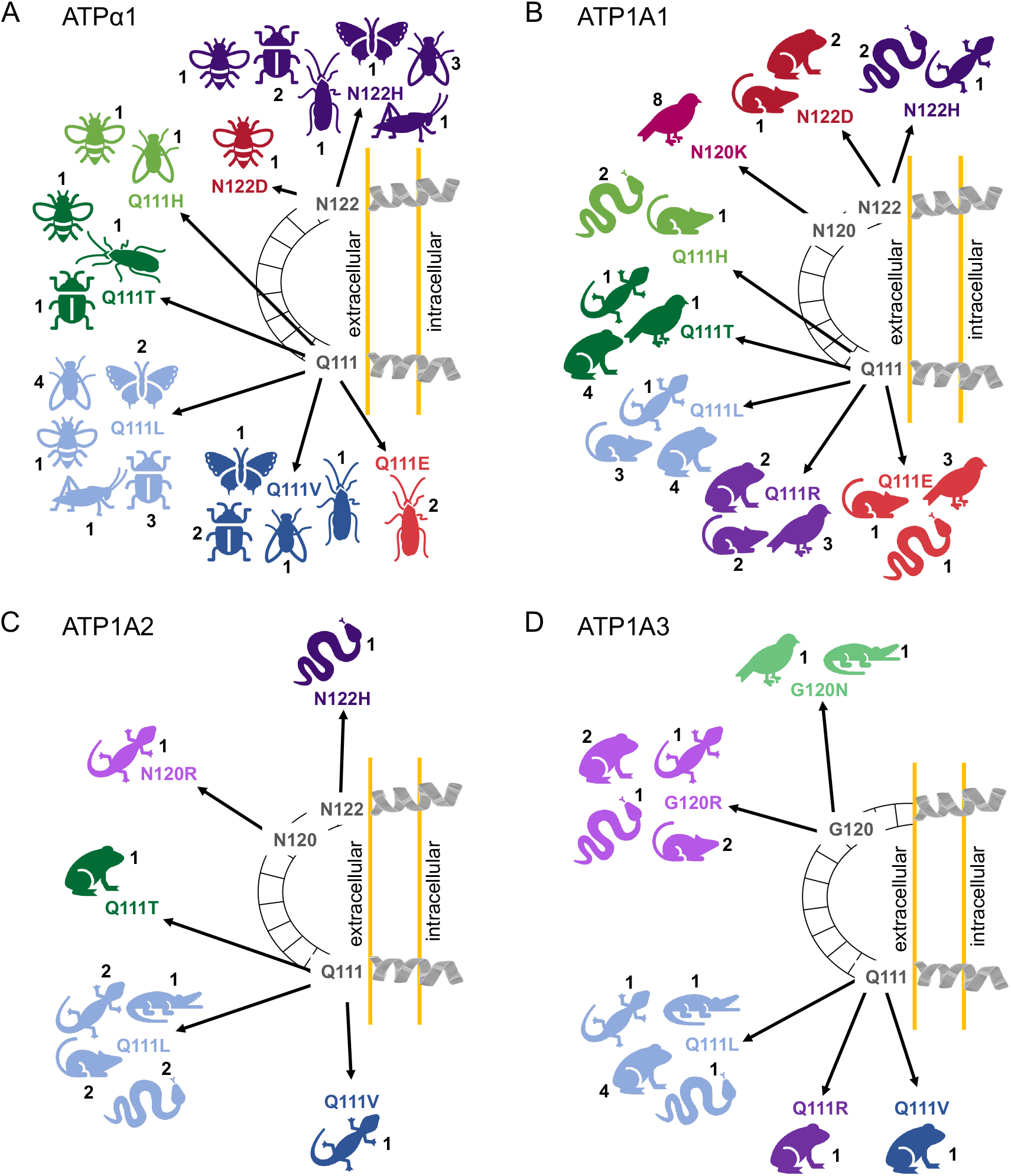
Parallel and divergent patterns of CTS-resistant substitutions across ATPα1 of insects and the shared ATP1A paralogs of tetrapods. Examples of convergence in ATPα1 across insects (A). Convergence in the **(B)** ATP1A1, **(C)** ATP1A2, and **(D)** ATP1A3 paralogs, respectively, across tetrapods. Numbers indicate the number of independent substitutions in each major clade depicted. For ATP1A3, resistance-conferring amino acid substitutions have been identified at site 120, and not 122. A full list of amino acid substitutions can be found in Supplementary Dataset 2 for tetrapods, and Taverner et al. [23] for insects.

**Figure 4.**
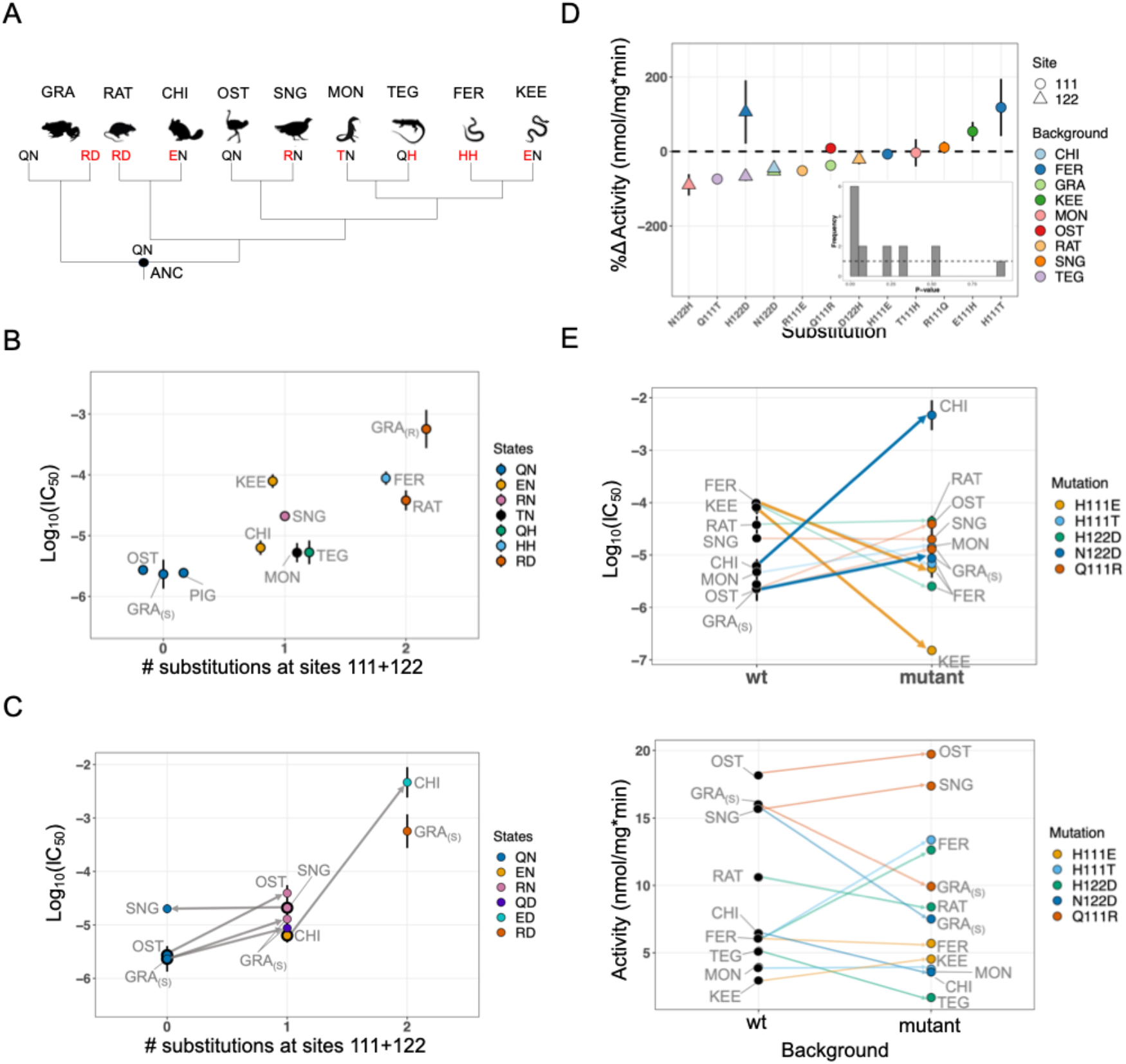
Functional properties of wild-type and engineered ATP1A1. **(A)** Cladogram relating the surveyed species. GRA: Grass Frog (*Leptodactylus*); RAT: Rat (*Rattus*); CHI: Chinchilla (*Chinchilla*); OST: Ostrich (*Struthio*); SNG: Sandgrouse (*Pterocles*); MON: Monitor lizard (*Varanus*); TEG: Tegu lizard (*Tupinambis*); FER: False fer-de-lance (*Xenodon*); KEE: Red-necked keelback snake (*Rhabdophis*). Two-letter codes underneath each avatar indicate native amino acid states at sites 111 and 122, respectively. Data for grass frog from Mohammadi et al. [25]. **(B)** Levels of CTS resistance (IC50) among wild-type enzymes. The x-axis distinguishes among ATP1A1 with 0, 1 or 2 derived states at sites 111 and 122. The subscripts S and R refer to the CTS-sensitive and CTS-resistant paralogs, respectively. **(C)** Effects on CTS resistance (IC_50_) of changing the number of substitutions at 111 or 122. Substitutions result in predictable changes to resistance except in the reversal R111Q in Sandgrouse (SNG). GRA_S_ represents Q111R+N122D on the sensitive paralog background. **(D)** Effects of single substitutions on Na,K-ATPase (NKA) activity. Each modified ATP1A1 is compared to the wild-type enzyme for that species. The inset shows the distribution of *t*-test p-values for all 15 substitutions, with the dotted line indicating the expectation. **(E)** Evidence for epistasis for CTS resistance (IC50, upper panel) and lack of such effects for enzyme activity (lower panel). Each line compares the same substitution (or the reverse substitution) tested on at least two backgrounds. Thicker lines correspond to substitutions with significant sequence-context dependent effects (Bonferroni-corrected ANOVA *p*-values < 0.05, Table S5).

To test for epistatic effects of common CTS-resistant substitutions at sites 111 and 122, we used site-directed mutagenesis to introduce 15 substitutions (nine at position 111 and six at position 122) in the wild-type ATP1A1 backgrounds of nine species (Fig. S6). The specific substitutions chosen were either phylogenetically broadly-distributed convergent substitutions and/or divergent substitutions that distinguish closely related clades of species. We expressed each of these 24 ATP1A1 constructs with an appropriate species-specific ATP1B1 protein (Table S3). For each recombinant NKA protein complex, we characterized its level of CTS resistance (IC_50_) and estimated enzyme activity as the rate of ATP hydrolysis in the absence of CTS (Table S4).

Among the 12 cases for which IC_50_ could be measured, substitutions had a 15-fold effect on average (Fig. 4C, Table S4) and were equally likely to increase or decrease IC_50_. To assess the background dependence of specific substitutions, we examined five cases in which a given substitution (e.g., E111H), or the reverse substitution (e.g., H111E), could be evaluated on two or more backgrounds. In the absence of intramolecular epistasis, the effect of a substitution on different backgrounds should remain unchanged and the magnitude of the effect of the reverse substitution should also be the same but with opposite sign. This analysis revealed substantial background dependence for IC_50_ in two of the five informative cases (Fig. 4E; Table S5). In one case, the N122D substitution resulted in a 200-fold larger increase in IC_50_ when added to the chinchilla (CHI) background compared to the grass frog (GRA) background (p=1.2e-3 by ANOVA). In the other case, the E111H substitution and the reverse substitution (H111E) produced effects in the same direction (reducing CTS-resistance) when added to different backgrounds (false fer-de-lance (FER) and red-necked keelback (KEE) snakes, respectively, p=1e-7 by ANOVA). Overall, these results suggest that the effect of a given substitution on IC_50_ can be strongly dependent on the background on which it occurs. The remaining three substitutions (H111T, Q111R and H122D) showed no significant change in the magnitude of the effect on IC_50_ when introduced into different species’ backgrounds. These results suggest that, while some substitutions can have strong background-dependent effects, strong intramolecular epistasis with respect to CTS resistance is not universal.

### Pleiotropic effects on NKA activity largely depend on states at a small set of sites

We next tested whether substitutions at sites 111 and 122 have pleiotropic effects on ATPase activity. Because ion transport across the membrane is a primary function of NKA and its disruption can have severe pathological effects [36], mutations that compromise this function are likely to be under strong purifying selection. As suggested by previous work [23–25], CTS-resistant substitutions at sites 111 and 122 can decrease enzyme activity. We evaluated the generality of these effects across a broader phylogenetic scale by comparing enzyme activity of the 15 mutant NKA proteins to their corresponding wild-type proteins.

Interestingly, the wild-type enzymes themselves exhibit substantial variation in activity, from 3-18 nmol/mg* min (*p*= 6e-7 by ANOVA, Fig 4E; Table S4). On average, substitutions at sites 111 and 122 on divergent orthologous protein backgrounds changed enzyme activity by 60% (mean of the absolute change; Fig 4D; Fig S4). In two cases, amino acid substitutions at position 122 (N122H and H122D) nearly inactivated lizard NKAs and, in one case, a substitution at position 111 (Q111T) resulted in low expression of the recombinant protein in the transfected cells (Fig S7; Fig. S8). A test of uniformity of pairwise t-test p-values across substitutions suggests a significant enrichment of low p-values (Fig 4D inset; p=2.5e-4, chi-squared test of uniformity). Thus, globally, this set of substitutions has significant effects on NKA activity, but they were surprisingly not more likely to decrease than increase activity (10 decrease : 5 increase, p>0.3, binomial test, Fig. 4D, Table S5).

We next asked to what extent pleiotropic effects of CTS resistant substitutions at positions 111 and 122 are dependent on genetic background. This question is motivated by recent studies in insects which revealed that deleterious pleiotropic effects of some resistance-conferring substitutions at sites 111 and 122 are background dependent [23,24]. Likewise, recent work on ATP1A1 of toad-eating grass frogs showed that effects of Q111R and N122D on NKA activity are also background dependent [25]. In contrast, among the five informative cases in which we compared the same substitution (or the reverse substitution) on two or more backgrounds, there is no evidence for background dependence (Fig 4E; Table S5). For example, N122D has similar effects on NKA activity in grass frog and chinchilla despite the substantial divergence between the species’ proteins (8.4% protein sequence divergence; Fig. 4D). Similarly, the effects of Q111R in ostrich or the reverse substitution R111Q in sandgrouse were not significantly different from the effect of Q111R in grass frog (7.5% and 8% protein sequence divergence, respectively).

To further examine the evidence for background dependence, we tested whether changes to the same amino acid state (regardless of starting state) at 111 and 122 produce different changes in NKA activity (e.g., R111E on the rat background versus H111E on the false fer-de-lance background). If epistasis is prevalent, involving a large number of sites, we expect that the absolute difference in effects of substitutions to a given amino acid state should increase with increasing sequence divergence of the wild-type ATP1A1 proteins. The 11 possible comparisons reveal substantial variation in the absolute difference of effects on protein activity, ranging from 8% to 190% (Table S7). Despite this, we found no relationship between the difference in the effect of substitutions to the same state and the extent of amino acid divergence between the orthologous proteins (Fig. 5A). This pattern suggests that, while pleiotropic effects may be pervasive and can be background dependent [23,25], these effects do not correlate with overall sequence divergence.

**Figure 5.**
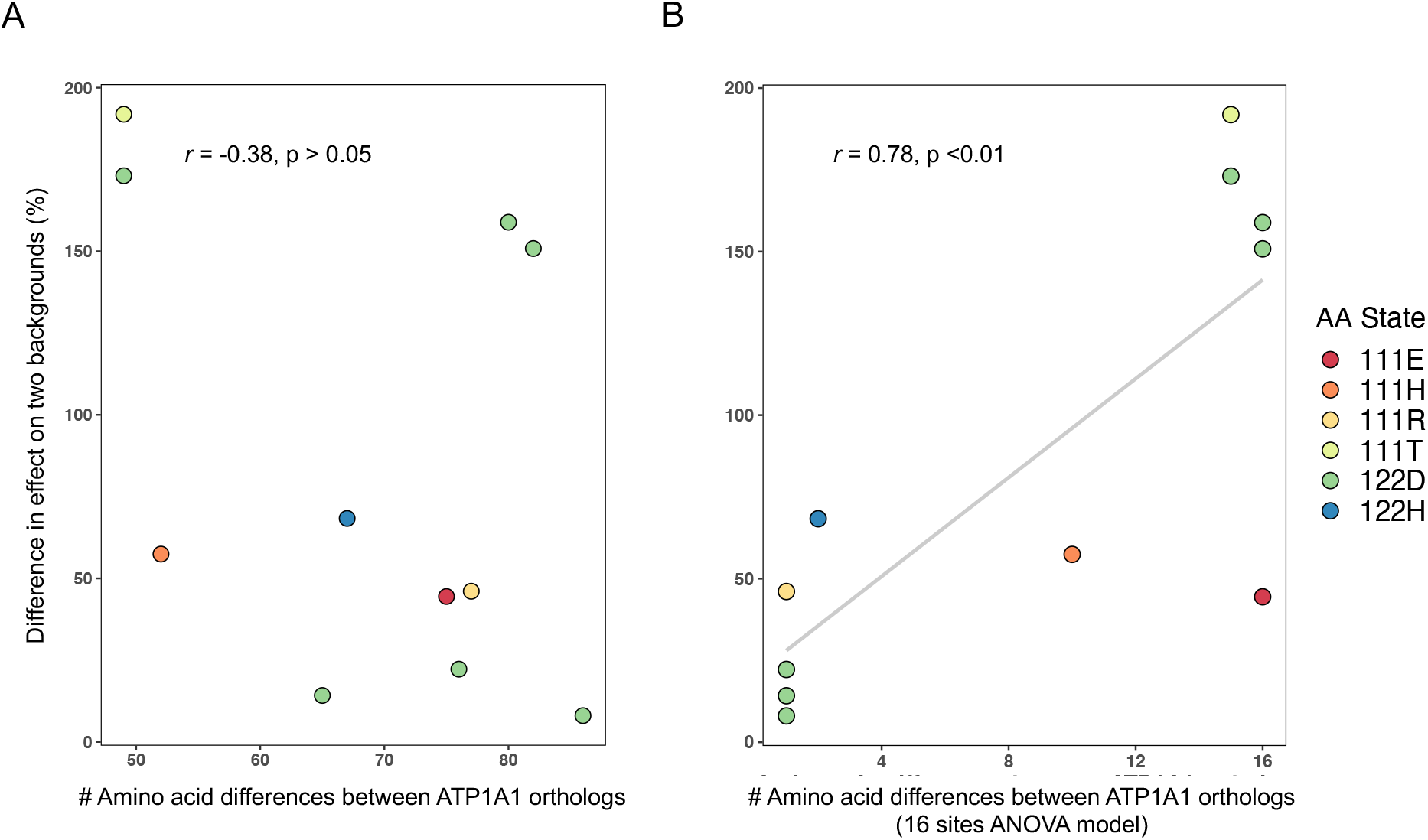
A small number of sites account for a large proportion of the differences in pleiotropic effects of the same substitution on divergent backgrounds. (A) The difference in effect size of a mutation to a given state on two different backgrounds as a function of divergence at all sites. Each point represents a comparison between the effect (% change in activity relative to the wild-type enzyme) of a given amino acid state (e.g.,122D) on two different genetic backgrounds. For example, the effect of 122D between chinchilla and false fer-de-lance is measured as | Δ% [chinchilla vs. chinchilla+N122D] minus the Δ% [false fer-de-lance vs. false fer-de-lance+H122D]|. Comparisons were measured as the difference between the two effects. The x-axis represents the number of amino acid differences between two wild-type ATP1A1 proteins (i.e., backgrounds) being compared. Assuming intramolecular epistasis for protein function is prevalent, a positive correlation is predicted. In total, 11 comparisons were possible, and no significant correlation is observed when considering divergence at all sites. (B) The difference in effect size of a mutation to a given state on two different backgrounds as a function of divergence at a subset of 16 sites with the largest effects on the difference in activity between two backgrounds. The p-value of the correlation was determined by permuting effects among constructs and generating a null distribution of correlations.

We hypothesized that background-dependent effects may instead depend on states at a small number of sites. If so, using total divergence may obscure a relationship between functional effects and divergence at these sites. To test this hypothesis, we used an analysis of variance to ask which variant sites across our functional constructs best accounted for differences in effects on different backgrounds (Methods and Fig. S5). Of 24 groups of variant sites (grouped as those with Pearson’s *r* > 0.8), we discovered two groups that included 16 of 113 total variant sites. These two groups of 16 sites jointly accounted for 78% of the variance among construct comparisons (p<0.004 by permutation). Further, in contrast to the pattern observed using all variant sites (Fig. 5A), we found a strong positive correlation between the difference in the effect of substitutions and the extent of divergence at these 16 sites (Fig. 5B; Pearson’s *r* =0.78, p=0.003 by permutation). This analysis strongly supports the notion that background-dependent effects depend on a circumscribed number of sites. While our resolution is limited (due to the limited number of genetic backgrounds in the experiments), we can say that 16 sites or fewer explain a large proportion of the differences in the effect of substitutions on different backgrounds.

### A global analysis of ATP1A sequences reveals further constraints on the evolution of CTS resistance

Since our functional experiments were necessarily limited in scope, we carried out a broad phylogenetic analysis to evaluate how well our findings align with global estimates of rates of convergence for the ATP1A family beyond ATP1A1 and beyond sites implicated in CTS resistance. Using a multisequence alignment of 831 ATP1A protein sequences, including the three ATP1A paralogs shared among tetrapods (i.e., amphibians, non-avian reptiles, birds, and mammals), we inferred a maximum likelihood phylogeny of the gene family (Fig. S1). We then used ancestral sequence reconstruction to infer the history of substitution events on all branches in the tree and counted the number of convergent amino acid substitutions per site along the entire protein (see Materials and Methods). Convergent substitutions are defined as substitutions on two branches at the same site resulting in the same amino acid state. Interestingly, we did not detect a correlation between the relative number of convergent substitutions with overall ATP1A divergence across the tree (Fig. S5). This result also held true when considering only substitutions to the key CTS-resistance sites 111 and 122 (Fig. S7).

To gain further insights into the factors that determine convergent evolution in ATP1A, we looked more closely at patterns of individual convergent substitutions at sites 111 and 122 by extracting each convergent substitution and visualizing its distribution along the sequence divergence axis (Fig. 6A). Under the expectation that rates of convergence should tend to decrease as a function of sequence divergence due to prevalent epistasis, the distribution of pairwise convergent events along the sequence divergence axis should be left-skewed, with a peak towards lower sequence divergence. In contrast to this expectation, the distribution is bimodal, with one peak at 0.33 and the other at 0.69 substitutions/site (Fig. 6B bottom panel). Convergent substitutions have occurred almost across the full range of protein divergence estimates. For example, if X is any starting state, the substitution X111R has occurred independently in 13 tetrapod lineages and X111L independently in 20 lineages. Both substitutions have a broad phylogenetic distribution, suggesting that their effects do not strongly depend on sequence states at a large number of sites throughout the protein. Interestingly, however, the distributions for X111H and X111E substitutions are relatively left-skewed, in line with epistasis for CTS resistance that we observed in experiments for H111E/E111H (Fig 4E).

**Figure 6.**
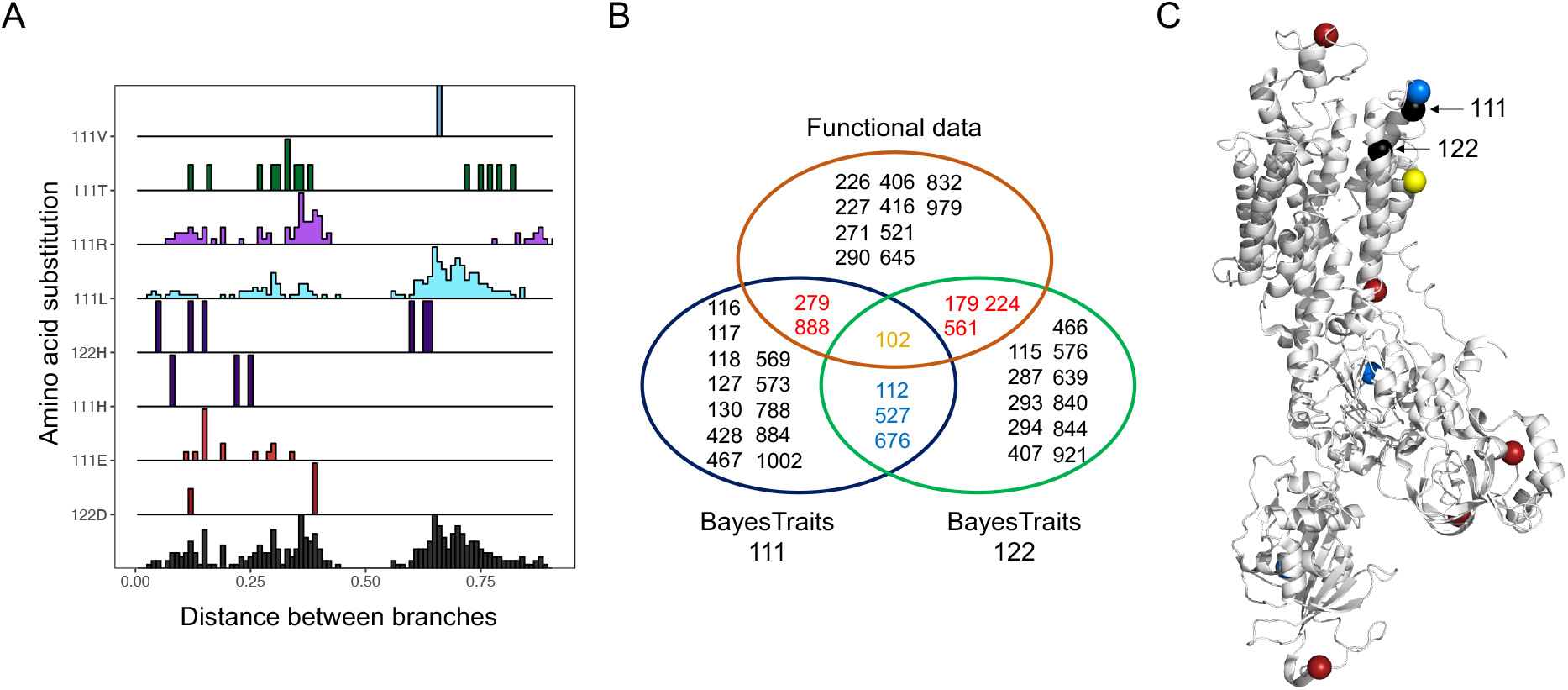
(A) Distribution of amino acid substitutions at sites 111 and 122 across all paralogs. For each derived amino acid state at sites 111 and 122, the histograms show the distribution of pairwise convergent events along the sequence divergence axis (expected number of substitutions per site). Substitutions are color coded as in Fig. 2. The histogram at the bottom shows the combined distribution of pairwise convergent events for both sites. (B) Intersection of 16 sites identified from functional data with the sites that most strongly correlated with substitutions at sites 111 and 122 Sites in the “functional data” group correspond to the 16 sites from the two groups identified by the ANOVA model (Tables S8 and S9). Sites in the BayesTraits analyses groups correspond to the top 5% sites with highest –log(P) association with 111 or 122, respectively. Overlaps between each group are larger than expected by chance: Functional∩BayesTraits111 = 3, *P* = 0.049; Functional∩ BayesTraits122 = 4, *P* = 0.007; BayesTraits111∩ BayesTraits122 = 4, *P* = 0.011. (C) Crystal structure of ATP1A1 (PDB 3B8E) showing sites color coded according to the intersections in panel B.

Using this same 831-sequence, 3-paralog alignment, we also asked which sites across all paralogs have substitution patterns that are most strongly correlated with those at 111 and 122 (Methods). We found 4 sites (102, 112, 527 and 676) that stand out as being in the top 5% of sites most strongly correlated with substitutions at both 111 and 122 (Fig. 6B), and this amount of overlap was larger than expected by chance. Further, the top 5% of sites most strongly correlated with site 111 also tend to be closer to 111 in atomic distance than expected (median distance to 111: 25.1 Å, p < 5e-4, bootstrap; Fig. 6C). Combining the results from our phylogenetic and functional analyses we identified a set of substitutions that are both strongly correlated with changes in 111 and 122 and account for a substantial proportion of the variance in background-dependent effects in our functional experiments (overlaps are larger than expected by chance; Fig 6B). For instance, despite being independently ascertained, site 102 is also among the most strongly predictive sites for background-dependent effects in our functional experiments, belonging to a group of 6 sites that explains 60% of variance (Fig. 5B and Fig. 6B). Together, our results suggest that proximate interactions involving a small number of sites, particularly site 111, are likely to be an important determinant of background-dependent effects.

## Discussion

Previous work has shown that the adaptive evolution of NKA-mediated CTS resistance in animals is constrained by pleiotropy and epistasis (i.e., background-dependence) [5,23–25,30]. Further, based on broad phylogenetic analysis of proteomes, our *a priori* expectation is that epistasis should represent a stronger constraint with increasing levels of divergence between species and ATP1A paralogs [20]. In light of these considerations, our extensive survey of the ATP1A gene family in tetrapods reveals two striking and seemingly contradictory patterns. The first is that some substitutions underlying CTS resistance in tetrapods are broadly distributed phylogenetically and even shared with insects (e.g., N122H is widespread among snakes and found in the monarch butterfly and other insects; see Fig. 3 for more examples). Patterns like these suggest that epistatic constraints have a limited role in the evolution of CTS resistance, as the same mutation can be favored on highly divergent genetic backgrounds. On the other hand, there is also substantial diversity in resistance-conferring states at sites 111 and 122, and some combinations of these substitutions appear to be phylogenetically restricted. For example, the CTS-resistant combination of Q111R+N122D has evolved multiple times in tetrapods but is absent in insects, whereas the CTS-resistant combination Q111V+N122H evolved multiple times in insects but is absent in tetrapods (Fig 3). Additionally, some substitutions also appear to be paralog-specific in tetrapods (Fig 3). These phylogenetic signatures suggest at least some role for epistasis as a source of contingency in the evolution of ATP1A-mediated CTS resistance in animals, i.e., that the fitness effects of substitutions depend on the order in which they occur. How can these disparate patterns be reconciled? To what extent do genetic background and contingency limit the evolution of CTS resistance in animals?

In our functional analysis of diverse ATP1A1 proteins, we find that derived substitutions at sites 111 and 122 have largely predictable effects on CTS resistance, with salient exceptions that tend to be in size rather than direction (Fig. 4). For example, Q111R contributes to CTS resistance on many species’ backgrounds, but not on that of sandgrouse (Fig. 4C and 4E). We also note that species with identical paired states at 111 and 122 can vary in CTS resistance by more than an order of magnitude (Fig. 4B). Together, these patterns point to background determinants of CTS resistance that may be additive rather than epistatic. Despite this, there are some substitutions that are widely distributed phylogenetically, such as N122D, that nonetheless do exhibit background-dependent effects on CTS resistance (Fig. 4C and 4E).

With respect to pleiotropic effects on NKA activity, our functional analysis of substitutions at sites 111 and 122 on diverse ATP1A1 backgrounds suggest that interactions between these substitutions and those backgrounds are largely additive. Specifically, we find that the severity of the effect of a particular CTS resistance substitution on NKA activity does not differ if added to protein backgrounds that are in the range of 49 to 86 amino acid substitutions away from the protein background on which that substitution naturally occurs (Fig. 5). In light of previous results demonstrating background-dependent effects of similar substitutions on NKA activity [23–25], our findings suggest that the extent of epistasis does not have a monotonic dependence on the extent of ATP1A1 divergence. Our findings further support increasing evidence that while epistasis is likely to be a pervasive feature of protein evolution, many mutational effects on structural and functional properties of proteins nonetheless seem to be additive (e.g., [37–39]).

We propose that our observations can be reconciled with previous results demonstrating epistatic constraints if epistasis with respect to protein function is confined to a small number of sites in the protein. If so, we might expect that the magnitude of epistasis may have little dependence on the extent of protein-wide ATP1A1 divergence but would instead be better predicted by divergence at a few key sites. In support of this view, Mohammadi et al. [25] showed that decreases in ATP1A1 enzyme activity due to substitutions Q111R and N122D can be rescued by 10 (or fewer) of the 19 amino acid differences distinguishing the backgrounds of CTS-resistant and sensitive ATP1A1 paralogs of grass frogs. Further, studies in *Drosophila melanogaster* [23,24] show that severe neural dysfunction associated with CTS resistance substitutions at sites 111 and 122 can largely be rescued with one additional substitution (A119S). Our study lends further support to this view by showing that one can identify a small group of sites (16 or fewer) that account for a large proportion of the variation in background-dependent effects on enzyme activity across proteins spanning the breath of the tetrapod phylogeny.

Phenotypes such as enzyme activity are not equivalent to organismal fitness and that there may be a nonlinear mapping between the two. The discussion above assumes that changes in enzyme activity are most likely detrimental to organismal fitness, but this need not be the case. We found that the activity of wild-type NKAs varies 6-fold among the species surveyed (Fig. 4E), suggesting that most species are either robust to changes in NKA activity or that changes have occurred in other genes (including other ATP1A paralogs) to compensate for changes in NKA activity associated with ATP1A1. Thus, either protein activity itself is not an important pleiotropic constraint on the evolution of CTS resistance of NKA, or constraint depends not just on the protein background but also on the broader genetic background of the organism (e.g., other interacting proteins; see [23]).

Our work highlights the utility of comparative functional work in understanding the nature of epistatic constraints on the evolution of novel protein functions. In this case-study of the evolution of CTS-resistant NKAs, we find that epistatic constraints are more likely to depend on divergence at a small number of key sites in the protein, likely in close proximity to site 111, rather than overall levels of protein divergence. The circumscribed nature of these constraints may account for the remarkable convergence of CTS-resistance substitutions observed among the NKAs of highly divergent species.

## Materials and Methods

### Sample collection and data sources

In order to carry out a comprehensive survey of vertebrate ATP1A paralogs (ATP1A1, ATP1A2 and ATP1A3), we collated a total of 831 protein sequences for this study (corresponding alignment can be found in Supplementary Dataset 1). Mammals possess a fourth paralog (ATP1A4) that is expressed predominantly in testes [40] that we did not consider here, although for completeness the protein sequences are provided in Supplementary Dataset 1 and the alignment of variant sites in Supplementary Dataset 2). The 831 sequences included RNA-seq data generated here for 27 species of non-avian reptiles (Table S1; PRJNA754197) to provide more information from some previously underrepresented lineages. These included field-caught and museum-archived specimens as well as animals purchased from commercial pet vendors. Purchased animals were processed following the procedures specified in the IACUC Protocol No. 2057-16 (Princeton University) and implemented by a research veterinarian at Princeton University. Wild-caught animals were collected under Colombian umbrella permit *resolución No*. 1177 granted by the *Autoridad Nacional de Licencias Ambientales* to the Universidad de los Andes and handled according to protocols approved by the Institutional Committee on the Care and Use of Laboratory Animals (abbreviated *CICUAL* in Spanish) of the Universidad de los Andes. In all cases, fresh tissues (brain, stomach, and muscle) were taken and preserved in RNAlater (Invitrogen) and stored at −80° C until used.

### Reconstruction of ATP1A paralogs

RNA-seq libraries were prepared either using TruSeq RNA Library Prep Kit v2 (Illumina) and sequenced on Illumina HiSeq2500 (Genomics Core Facility, Princeton, NJ, USA) or using NEBNext Ultra RNA Library Preparation Lit (NEB) and sequenced on Illumina HiSeq4000 (Genewiz, South Plainfield, NJ, USA) (Table S2). All raw RNA-seq data generated for this study have been deposited in the National Center for Biotechnology Information (NCBI) Sequence Read Archive (SRA) under Bioproject PRJNA754197. Together with SRA datasets downloaded from public database, reads were trimmed to Phred quality ≥ 20 and length ≥ 20 and then assembled *de novo* using Trinity v2.2.0 [41]. Sequences of ATP1A paralogs 1, 2 and 3 were pulled out with BLAST searches (blast-v2.26), individually curated, and then aligned using ClustalW. Complete alignments of ATP1/2/3 can be found in Supplementary Dataset 1.

### Statistical phylogenetic analyses

We use the standard sheep (*Ovis aries*) numbering system to match previous literature; this number corresponds to the Uniprot:P04074 sequence minus 5 residues from the 5’ end. Protein sequences from ATP1A1 (N=429), ATP1A2 (N=197) and ATP1A3 (N=205) including main tetrapod classes (amphibians, non-avian reptiles, birds, and mammals) plus lungfish and coelacanth as outgroups were aligned using ClustalW with default parameters. The optimal parameters for phylogenetic reconstruction were taken from the best-fit amino acid substitution model based on Akaike Information Criterion (AIC) as implemented in ModelTest-NG v.0.1.5 [42], and was inferred to be JTT+G4+F. An initial phylogeny was inferred using RAxML HPC v.8 [43] under the JTT+GAMMA model with empirical amino acid frequencies. Branch lengths and node support (aLRS) were further refined using PhyML v.3.1 [44] with empirical amino acid frequencies and maximum likelihood estimates of rate heterogeneity parameters, I and Γ. Phylogeny visualization and mapping of character states for each paralog was done using the R package ggtree [45].

### Ancestral sequence reconstruction and convergence calculations

Ancestral sequence reconstruction was performed in PAML using codeml [46] under the JTT+G4+F substitution model. Ancestral sequences from all nodes in the ATP1A phylogeny were combined with extant sequences to produce a 1040 amino-acid multiple sequence alignment of 1,660 ATP1A proteins (831 extant species and 829 inferred ancestral sequences; Fig. S1). For each branch in the tree, we determined the occurrence of substitutions by using the ancestral and derived amino acid states at each site using only states with posterior probability (PP) > 0.8. All branch pairs were compared, except sister branches and ancestor-descendent pairs [20,21]. When comparing substitutions on two distinct branches at the same site, substitutions to the same amino acid state were counted as convergences, while substitutions away from a common amino acid were counted as divergences. We excluded a putative 30 amino acid-long alternatively-spliced region (positions 810-840). For each pairwise comparison, we calculated the proportion of observed convergent events per branch as (number of convergences +1) / (number of divergences +1). The line describing the trend was calculated as a running average with a window size of 0.05 substitutions/site. 95% confidence intervals were estimated based on 100 random samples of pairwise branch comparisons for each window. To determine whether convergence at sites 111 or 122 decreases with sequence divergence, we encoded convergence events as “1” and divergence events as “0” for all pairwise sequence comparisons and used a logistic regression to test for the correlation between molecular convergence (0 or 1) and genetic distance (Fig. S4).

### Identifying correlated substitutions

We used BayesTraits [47] to detect sites across the ATP1A phylogeny that exhibit correlated evolution with sites 111 and 122. Using the reconstructed ancestral sequences for each paralogous clade (i.e., the most recent common ancestor (MRCA) of ATP1A1, the MRCA of ATP1A2, and the MRCA of ATP1A3), we coded each amino acid state among extant sequences of the multi-species alignment into ancestral ‘0’ and derived ‘1’ states, and used these plus the phylogeny with estimated branch lengths as inputs for BayesTraits. BayesTraits fits a continuous-time Markov model to estimate transition rates between discrete, binary traits and estimates the best fitting model describing their joint evolution on a phylogeny. Specifically, we tested whether the rate of evolution at sites 111 and 122 was dependent on all other variant sites (see below). We excluded singleton sites and sites with more than 80% gaps, as these sites would be of little information, resulting in an analysis of 417 variant sites.

We tested two sets of models, hereafter referred as base and restricted models, each of which has a null independent and an alternative dependent model. The null independent model assumes that the two sites evolve independently, and the alternative dependent model assumes that the sites are correlated such that the change at one site is dependent on the state at the other site. Because the null model is a general form of the alternative model, both models can be compared under a likelihood ratio test (LRT) with degrees of freedom (df) equal to the difference in the number of parameters between models. The base and restricted sets of models differ on the presence of restricted parameters for certain transition rates, and consequently differ in the number of df. In the base models, the null model has four rate parameters describing each possible independent change of state at each site; the alternative model has eight rate parameters describing all possible changes at each site dependent on the state at the other site (LRT with df=4). For the restricted models, we set the rates of transition to the ancestral state to zero [23] as the median branch length of the tree of 0.002351 substitutions per site makes it unlikely for a site to change twice or back to the ancestral state. After these restrictions, the null independent model had four transition parameters. To test for dependence, we imposed two additional restrictions to the model: one forcing the transition rate at site one to be fixed regardless of the state of site 2 (q13=q24), and a second forcing the transition rate at site 2 to be fixed regardless of the state of site 1 (q12=q34) [23]. This effectively tests whether the transition rate is affected by the state of either site and leaves the model with only two transition parameters (LRT with df=2). To run the analysis, the phylogeny branch lengths were scaled using BayesTraits to have a mean length of 0.1, and to increase the chance of finding the true maximum likelihood, we set MLTries to 250.

### Protein structure analysis

To test for spatial clustering of sites showing statistical signatures of coevolution with site 111 or 122, we used a custom Python script (available on request) and the Bio.PDB’s module. We used the crystal structure of Na,K-ATPase (PDB: 3b8e) to estimate distances (in Angstroms) between the alpha carbon of site 111 or 122 to the alpha carbon of all other variable sites in the alignment. We calculated the median distance of the top 5% of variable sites with the strongest signature of correlated evolution with each focal site, 111 or 122 (from BayesTraits output). We estimated the p-value using 1000 random samples of 5% of variable sites and calculating the proportion of times the median value was less than or equal to the observed value.

### Construction of expression vectors

ATP1A1 and ATP1B1 wild-type sequences for the eight selected tetrapod species (Fig. 4) were synthesized by Invitrogen™ GeneArt. The *β*1-subunit genes were inserted into pFastBac Dual expression vectors (Life Technologies) at the p10 promoter with XhoI and PaeI (FastDigest Thermo Scientific™) and then control sequenced. The α1-subunit genes were inserted at the PH promoter of vectors already containing the corresponding *β*1-subunit proteins using In-Fusion® HD Cloning Kit (Takara Bio, USA Inc.) and control sequenced. All resulting vectors had the α1-subunit gene under the control of the PH promoter and a *β*1-subunit gene under the p10 promoter. The resulting eight vectors were then subjected to site-directed mutagenesis (QuickChange II XL Kit; Agilent Technologies, La Jolla, CA, USA) to introduce the codons of interest. In total, 21 vectors were produced (Table S3).

### Generation of recombinant viruses and transfection into Sf9 cells

*Escherichia coli* DH10bac cells harboring the baculovirus genome (bacmid) and a transposition helper vector (Life Technologies) were transformed according to the manufacturer’s protocol with expression vectors containing the different gene constructs. Recombinant bacmids were selected through PCR screening, grown, and isolated. Subsequently, Sf9 cells (4 × 10^5^ cells* ml) in 2 ml of Insect-Xpress medium (Lonza, Walkersville, MD, USA) were transfected with recombinant bacmids using Cellfectin reagent (Life Technologies). After a three-day incubation period, recombinant baculoviruses were isolated (P1) and used to infect fresh Sf9 cells (1.2 × 10^6^ cells* ml) in 10 ml of Insect-Xpress medium (Lonza, Walkersville, MD, USA) with 15 mg/ml gentamycin (Roth, Karlsruhe, Germany) at a multiplicity of infection of 0.1. Five days after infection, the amplified viruses were harvested (P2 stock).

### Preparation of Sf9 membranes

For production of recombinant NKA, Sf9 cells were infected with the P2 viral stock at a multiplicity of infection of 10^3^. The cells (1.6 × 10^6^ cells* ml) were grown in 50 ml of Insect-Xpress medium (Lonza, Walkersville, MD, USA) with 15 mg/ml gentamycin (Roth, Karlsruhe, Germany) at 27°C in 500 ml flasks (35). After 3 days, Sf9 cells were harvested by centrifugation at 20,000 x g for 10 min. The cells were stored at −80 °C and then resuspended at 0 °C in 15 ml of homogenization buffer (0.25 M sucrose, 2 mM EDTA, and 25 mM HEPES/Tris; pH 7.0). The resuspended cells were sonicated at 60 W (Bandelin Electronic Company, Berlin, Germany) for three 45 s intervals at 0 °C. The cell suspension was then subjected to centrifugation for 30 min at 10,000 x g (J2-21 centrifuge, Beckmann-Coulter, Krefeld, Germany). The supernatant was collected and further centrifuged for 60 m at 100,000 x g at 4 °C (Ultra-Centrifuge L-80, Beckmann-Coulter) to pellet the cell membranes. The pelleted membranes were washed once and resuspended in ROTIPURAN® p.a., ACS water (Roth) and stored at −20 °C. Protein concentrations were determined by Bradford assays using bovine serum albumin as a standard. Three biological replicates were produced for each NKA construct.

### Verification by SDS-PAGE/western blotting

For each biological replicate, 10 μg of protein were solubilized in 4x SDS-polyacrylamide gel electrophoresis sample buffer and separated on SDS gels containing 10% acrylamide. Subsequently, they were blotted on nitrocellulose membrane (HP42.1, Roth). To block non-specific binding sites after blotting, the membrane was incubated with 5% dried milk in TBS-Tween 20 for 1 h. After blocking, the membranes were incubated overnight at 4 °C with the primary monoclonal antibody α5 (Developmental Studies Hybridoma Bank, University of Iowa, Iowa City, IA, USA). Since only membrane proteins were isolated from transfected cells, detection of the α subunit also indicates the presence of the β subunit. The primary antibody was detected using a goat-anti-mouse secondary antibody conjugated with horseradish peroxidase (Dianova, Hamburg, Germany). The staining of the precipitated polypeptide-antibody complexes was performed by addition of 60 mg 4-chloro-1 naphtol (Sigma-Aldrich, Taufkirchen, Germany) in 20 ml ice-cold methanol to 100 ml phosphate buffered saline (PBS) containing 60 μl 30% H_2_O_2_. See Fig. S8.

### Ouabain inhibition assay

To determine the sensitivity of each NKA construct against cardiotonic steroids (CTS), we used the water-soluble cardiac glycoside, ouabain (Acrō s Organics), as our representative CTS. 100 ug of each protein was pipetted into each well in a nine-well row on a 96-well microplate (Fisherbrand) containing stabilizing buffers (see buffer formulas in [48]). Each well in the nine-well row was exposed to exponentially decreasing concentrations of ouabain (10^−3^ M, 10^−4^ M, 10^−5^ M, 10^−6^ M, 10^−7^ M, 10^−8^ M, dissolved in distilled H_2_O), plus distilled water only (experimental control), and a combination of an inhibition buffer lacking KCl and 10^−2^ M ouabain to measure background protein activity [48]. The proteins were incubated at 37°C and 200 rpms for 10 minutes on a microplate shaker (Quantifoil Instruments, Jena, Germany). Next, ATP (Sigma Aldrich) was added to each well and the proteins were incubated again at 37°C and 200 rpms for 20 minutes. The activity of NKA following ouabain exposure was determined by quantification of inorganic phosphate (Pi) released from enzymatically hydrolyzed ATP. Reaction Pi levels were measured according to the procedure described in Taussky and Shorr [49] (see Petschenka et al. [48]). All assays were run in duplicate and the average of the two technical replicates was used for subsequent statistical analyses. Absorbance for each well was measured at 650 nm with a plate absorbance reader (BioRad Model 680 spectrophotometer and software package). See Table S4.

### ATP hydrolysis assay

To determine the functional efficiency of different NKA constructs, we calculated the amount of Pi hydrolyzed from ATP per mg of protein per minute. The measurements (the mean of two technical replicates) were obtained from the same assay as described above. In brief, absorbance from the experimental control reactions, in which 100 μg of protein was incubated without any inhibiting factors (i.e., ouabain or buffer excluding KCl), were measured and translated to mM Pi from a standard curve that was run in parallel (1.2 mM Pi, 1 mM Pi, 0.8 mM Pi, 0.6 mM Pi, 0.4 mM Pi, 0.2 mM Pi, 0 mM Pi). See Table S4.

### Statistical analyses of functional data

ATPase activity in the presence and absence of the CTS ouabain was measured following Petschenka et al. [48]. Background phosphate absorbance levels from reactions with inhibiting factors were used to calibrate phosphate absorbance. For ouabain sensitivity measurements, these calibrated absorbance values were converted to percentage non-inhibited NKA activity based on measurements from the control wells (as above). For each of the 3 biological replicates, log10 IC_50_ values were estimated using a four-parameter logistic curve, with the top asymptote set to 100 and the bottom asymptote set to zero, using the nlsLM function of the minipack.lm library in R [25]. To measure baseline recombinant protein activity, the calculated Pi concentrations of 100 μg of protein assayed in the absence of ouabain were converted to nmol Pi/mg protein/min. We used paired *t*-tests with Bonferroni corrections to identify significant differences between constructs with and without engineered substitutions. We used a two-way ANOVA to test for background dependence of substitutions (i.e., interaction between background and amino acid substitution) with respect to ouabain resistance (log10 IC_50_) and protein activity. Specifically, we tested whether the effects of a substitution X->Y are equal on different backgrounds (null hypothesis: X->Y (background 1) = X->Y (background 2)). We further assumed that the effects of a substitution X->Y should match that of Y->X. All statistical analyses were implemented in R. Data were plotted using the ggplot2 package in R.

Additionally, we evaluated the relationship between the effect of substitutions to a given amino acid state and the extent of sequence divergence between the protein backgrounds on which these substitutions were tested. To do this, we first calculated the effect of introducing a derived amino acid state as the percent change in protein activity relative to the wild-type protein. For example, the effect of the mutation N122D in Chinchilla (CHI) is

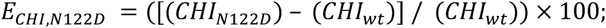

We then calculated the absolute difference between effects of substitutions to the same amino acid on two different backgrounds. For example, the difference (Δ) in the effect of 122D when introduced to the Chinchilla (CHI, mammal) and false fer-de-lance (FER, snake) proteins is

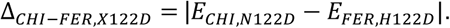

These calculations were possible for 11 pairwise comparisons (4 for site 111 and 7 for site 122; Table S7). We then evaluated the relationship between the estimated differences in the effects of substitutions to a given state versus the extent of protein sequence divergence (number of amino acid differences) between wild-type backgrounds.

To identify variant sites that most strongly predicted background-dependent effects in our data, we employed a site-by-site ANOVA analysis. For each of the eleven pairwise comparisons (e.g., Δ_*CHI−FER, X122D*_) each variant site was encoded as ‘0’ or ‘1’ if the wild-type sequences had the same or different amino acid state, respectively. This binarized per-site divergence (0 or 1) was used as the dependent variable in the ANOVA, with sites 111 or 122 (the mutated sites) as a covariate (Fig. S5A). For each of the 113 variant sites among the eight wild-type proteins, we then estimated that site’s marginal variance explained.

Given the limited number of wild-type backgrounds (8) relative to the number of sites (113), and use of constructs in multiple comparisons, strong correlations occur some variant sites (Fig. S5B). We thus grouped sites according to how they partition the divergence in experimental pairwise sequence comparisons. Grouping sites with Pearson’s r >0.8 results in 24 groups. Using one representative site per group, we then fitted nested ANOVA models to determine how much of variation in the Δ is explained by adding an additional group of sites, adding groups in the order of largest (group 1) to smallest (group 24) amount of variance explained. Using Likelihood Ratio Tests (LRTs) and Akaike’s Information Criteria (AIC), we identified the best model as the one including only the first two groups, which represent a total of 16 sites (14 sites in group 1 and two sites in group 2; Fig. S5C and Table S9). These 16 sites account for 78% of the variance (ANOVA R^2^). Since groups 1 and 2 were ascertained as those accounting for the largest proportion of the variance, we established the significance of this observation by permutation. Specifically, we performed 10,000 permutations of the experimental pairwise Δ across construct comparisons and repeated the procedure that was applied to the observed data to obtain a null distribution of R^2^ values. The p-value is estimated as the probability of finding two groups of sites that explain R^2^ ≥0.78. We further evaluated the extent of correlation (estimated as Pearson’s r) between Δ and sequence divergence at the 16 sites identified above. Similarly, to test for the significance of our regression model between Δ and divergence, we estimated the p-value as the probability of observing a Pearson’s r of 0.78 (or R^2^ of 0.61) or larger based on 10,000 permuted samples (permuting effects, Δ, across construct comparisons). To determine the robustness of our results to the grouping criteria, we did the same analyses using a higher cutoff of Pearson’s r > 0.99 (Table S8).

## Acknowledgments

We thank G. Sella and M. Przeworski for helpful discussions, and M. Przeworski for critical reading of an early draft of this paper. We thank C. Natarajan, P. Kowalski, M. Winter, and V. Wagschal for assistance in the laboratory, and D.A. Gómez-Sánchez for assistance in the field. We thank J. Oaks for providing tissue from ring-necked snake. We thank the Vice-president’s Office for Research and Creation of the Univerisdad de los Andes for help with permits. This study was funded by grants from the National Institutes of Health to PA (R01-GM115523), JFS (R01-HL087216) and SM (F32– HL149172), the National Science Foundation (OIA-1736249) to JFS, the Deutsche Forschungsgemeinschaft (Do 517/10-1) to SD, and the Alexander von Humboldt Foundation to SM.

## Notes

### Competing Interest Statement

The authors have declared no competing interest.

