## Supplemental Information for "Constraints on the evolution of toxin-resistant Na,K-ATPases have limited dependence on sequence divergence"

**This PDF file includes:**

Figures S1 to S8
Tables S1 to S6
SI References

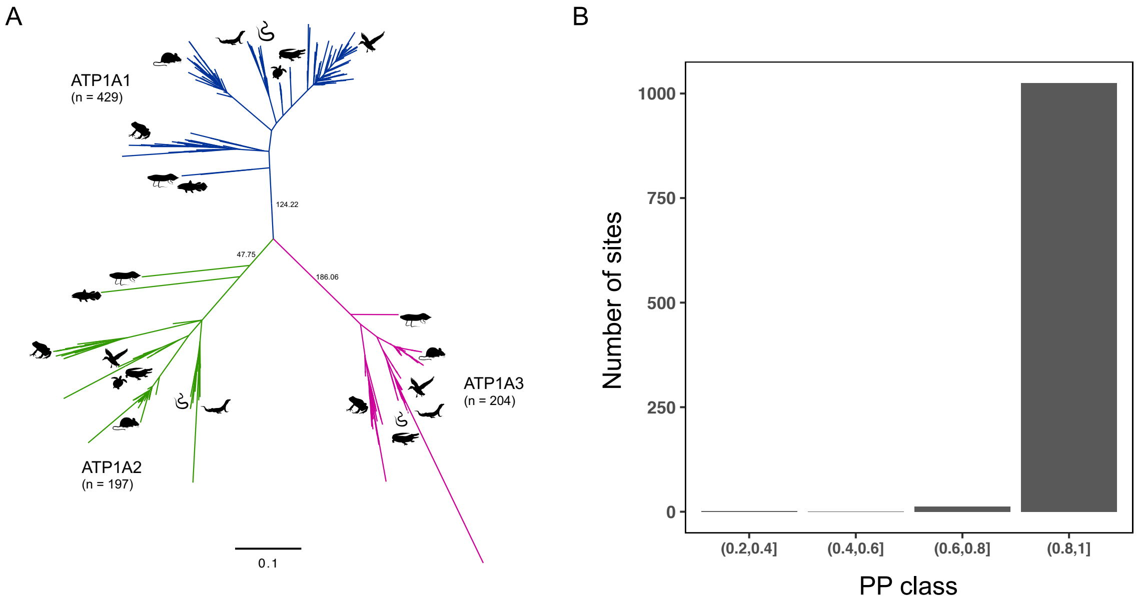


**Fig. S1.** (A) Maximum likelihood phylogeny of the ATP1A protein family. Colored clades correspond to each paralog, showing branch supports as approximate-likelihood ratio statistic (aLRS). Bottom scale bar shows the expected number of substitutions per site. (B) Ancestral sequence reconstruction for the ATP1A protein family. Barplot showing the counts of sites by posterior probability (PP) class. Each site is assigned to a class based on the mean PP across 829 ancestral sequences.

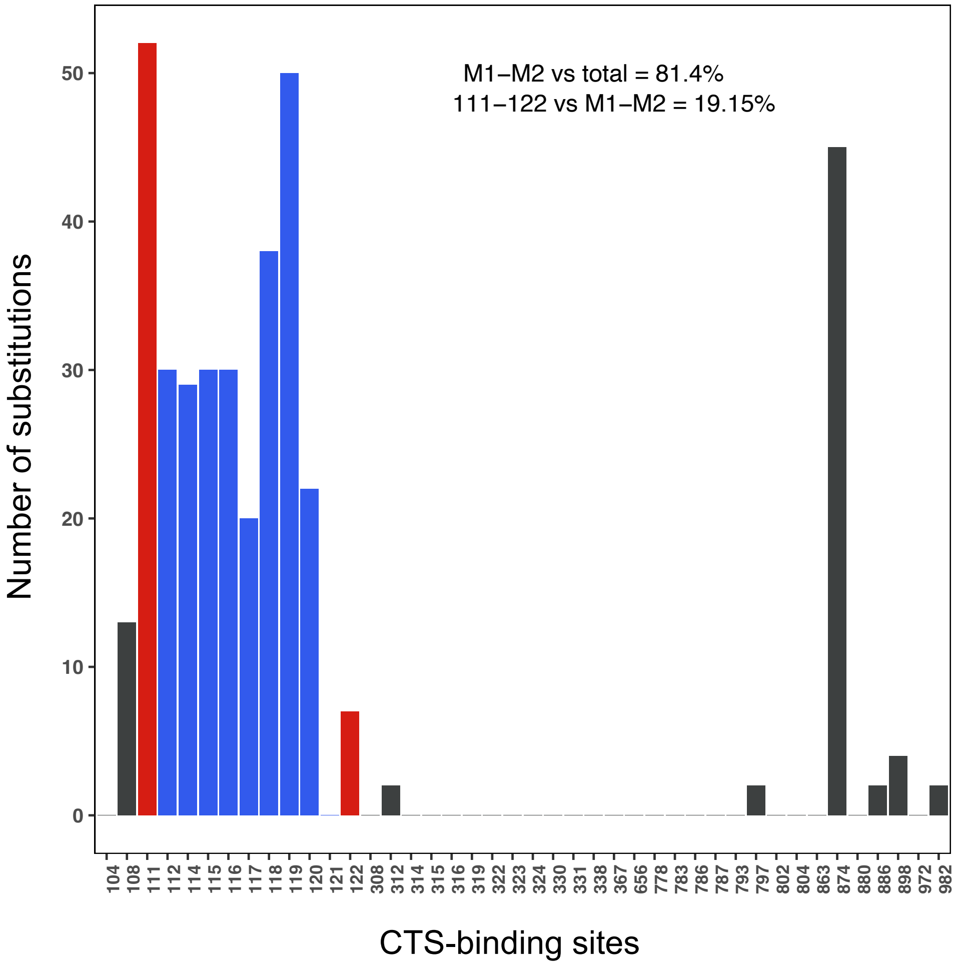


**Fig. S2.** Phylogenetic count of substitutions that occurred on 42 sites known to be implicated in CTS resistance across the complete ATP1A protein family (Fig S1A). Sites in the H1-H2 loop (M1-M2) account for 81% of the total number of substitutions that occurred in the 42 sites; sites 111 and 122 account for 19% of the substitutions observed in the H1-H2 loop. Sites are grouped as follows: 111-122 (red), H1-H2 extracellular loop (blue), and other sites (gray).

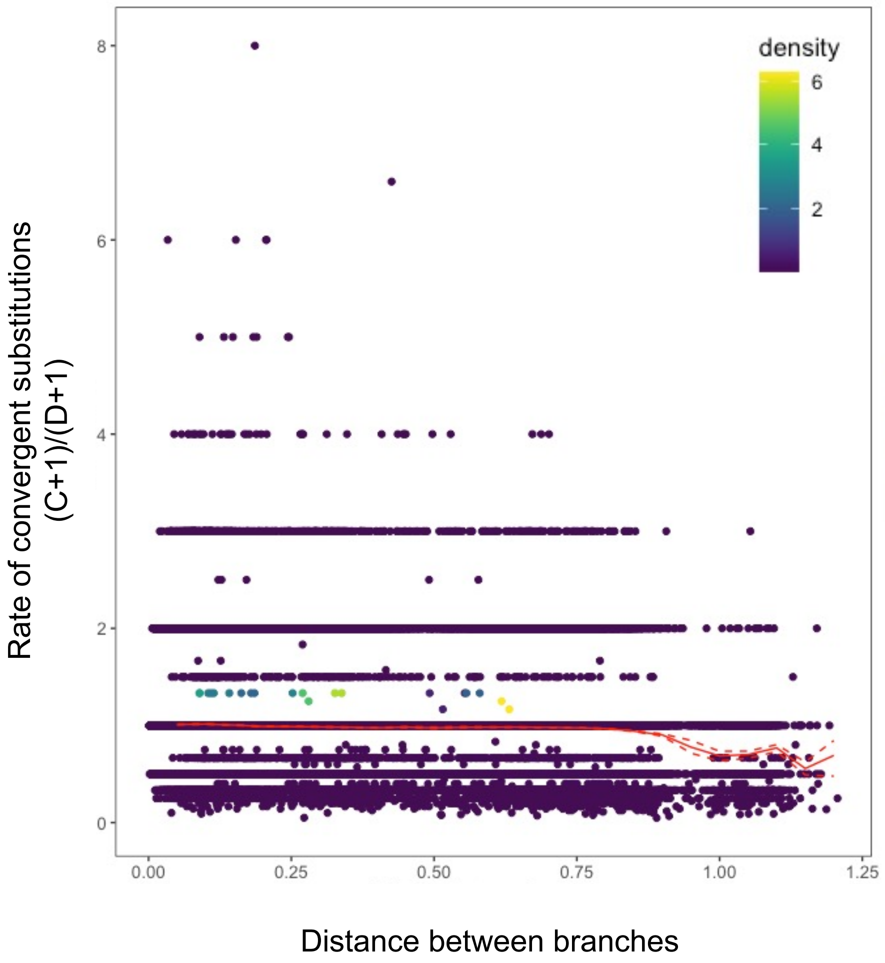


**Figure S3.** Rate of convergence across ATP1A sequences as a function of increasing sequence divergence. Change in the rate of convergence (protein wide) over time for the ATP1A protein family. The proportion of convergent (C) over divergent (D) substitutions along the entire protein sequence was estimated for all pairs of branches in the ATP1A phylogeny, except for sister branches or ancestor-descendant pairs. Color scale shows the density of dots for both axes. The distance between branches corresponds to the expected number of amino acid substitutions per site between protein pairs being compared (under the JTT+G4+F model). The red line shows a running average with a window size of 0.05 substitutions/site. Dashed lines show the 95% confidence interval based on 100 bootstrap replicates per window

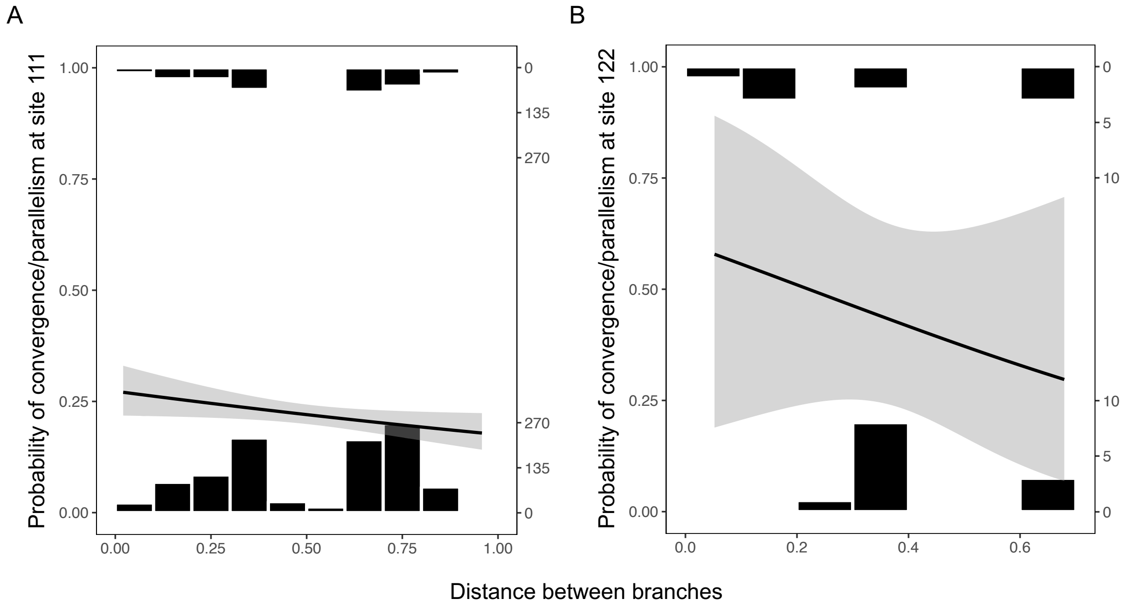


**Fig. S4.** Probability of convergence and parallelism at sites (A) 111 and (B) 122 as a function of the distance between branches (i.e. the sum of internal branch lengths between two proteins). Histograms show the phylogenetic distribution of events where a substitution at site 111 or 122 on two branches resulted in the same amino acid state (top) or to a different state (bottom), with substitution counts indicated on the righthand vertical axis. Site 111: log-odds -0.5673, P=0.038; site 122: log-odds -1.8788, P>0.1.

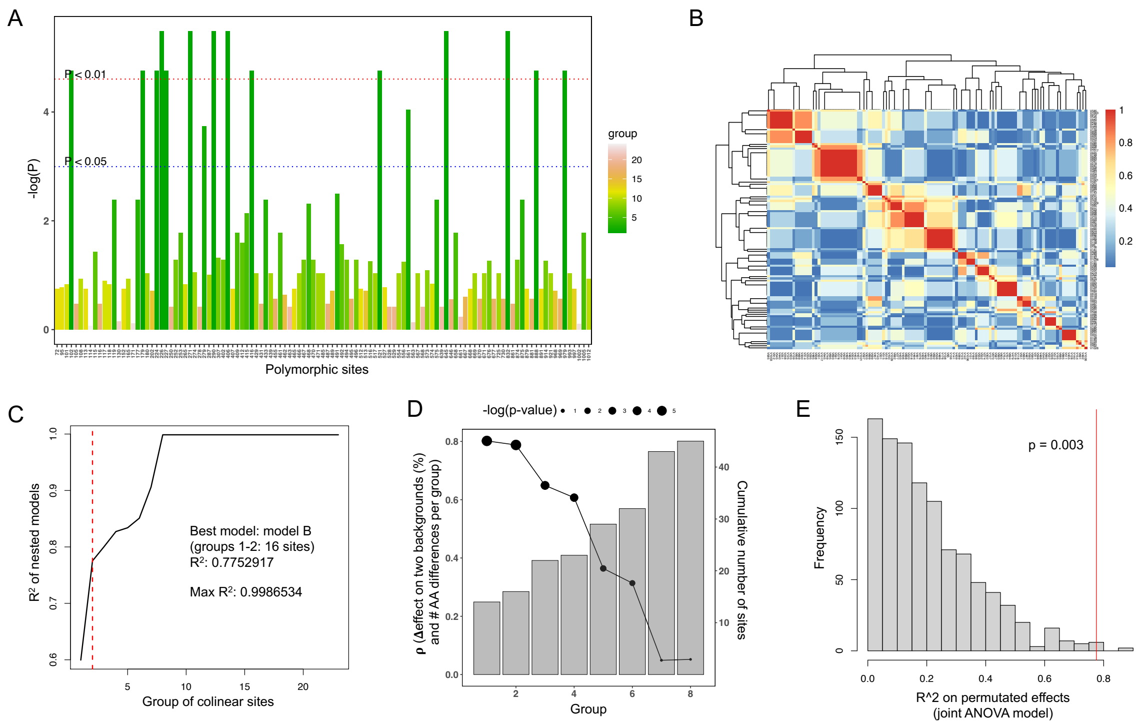


**Figure S5. A)** "Manhattan plot" showing the significance of the extent of variance in measured effects explained by divergence at each variant site. Note that sites with high –log(P) values are not independent. **B)** Pearson's correlation coefficients (r) among 113 variant sites distinguishing the 8 wild-type constructs (i.e. backgrounds). We used a cutoff of $|r| > 0.8$ to define groups of highly correlated sites for model analyses. **C)** Proportion of variation ($R^2$) that is explained by increasingly nested ANOVA models using one representative site per group of colinear sites. Red dashed line shows the best model (model B: groups 1+2) based on a likelihood ratio test (LRT) and Akaike Information Criteria (AIC). **D)** Pearson's r (% Δeffect on two backgrounds vs. number of amino acid differences per groups) decreases monotonically with the number of sites included (bars – right Y-axis). The correlations remain significant ($P<0.05$) up to the model D (groups 1+2+3+4) which includes a total of 23 sites. **E)** Null distribution of $R^2$ values for the best model (model B: groups 1+2) based on permutated effects to the same derived amino acid state among sites. The red line shows the observed $R^2$ for the best model. The P-value was calculated as the probability of observing $R^2_{null} >= R^2_{obs}$.

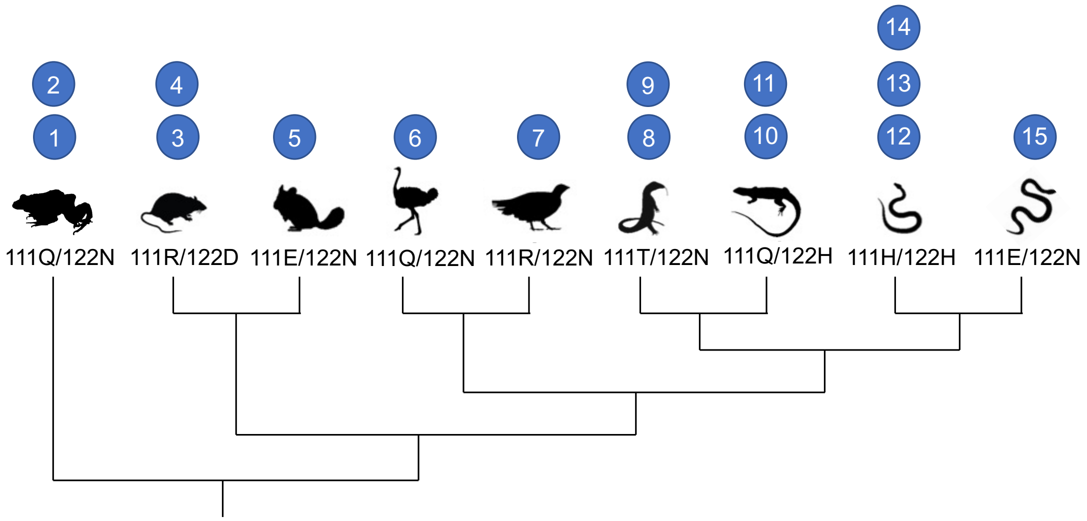


1. Q111R, grass frog + 111R found in frogs, rodents, birds, crocodilians, and turtles
2. N122D, grass frog + 122D found in frogs and rodents
3. R111E, rat + 111E found in rodents, birds, and snakes
4. D122H, rat + 122H found in lizards and snakes
5. N122D, chinchilla + 122D found in frogs and rodents
6. Q111R, ostrich + 111R found in frogs, rodents, birds, crocodilians, and turtles
7. R111Q, yellow-throated sandgrouse + 111Q found in all animals
8. T111H, Savannah monitor + 111H found in mammals and snakes
9. N122H, Savannah monitor + 122H found in lizards and snakes
10. Q111T, black tegu + 111T found in frogs, birds, and lizards
11. H122D, black tegu + 122D found in frogs and rodents
12. H122D, false fer-de-lance + 122D found in frogs and rodents
13. H111T, false fer-de-lance + 111T found in frogs, birds, and lizards
14. H111E, false fer-de-lance + 111E found in rodents, birds, and snakes
15. E111H, red-necked keelback + 111H found in mammals and snakes

**Fig. S6.** Diagram illustrating the experimental design of recombinant Na,K-ATPase proteins with amino acid substitutions at positions 111 and 122.

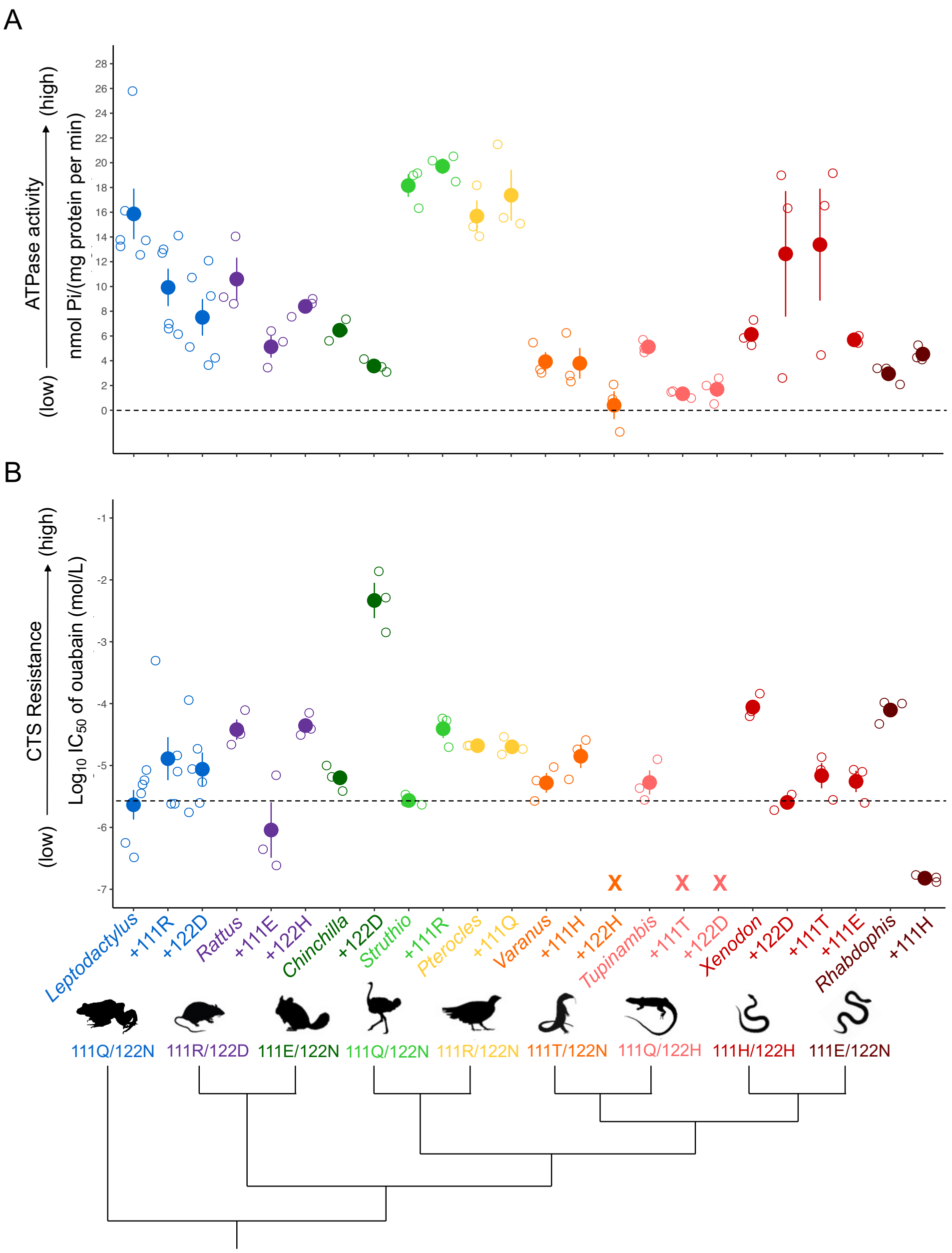


**Fig. S7.** Joint functional properties of 24 engineered Na,K-ATPases (NKAs) from eight vertebrate species. Proteins are color-coded according to the species from which they were engineered. Wildtype amino acid states at position 111 and 122 for each species and phylogenetic relationships are denoted at the bottom. For each species, from left to right, functional data for the wildtype protein are first shown (denoted by the genus name), followed by the mutant proteins (denoted by +[mutation]). Panel A shows mean ± SEM ATP hydrolysis activity (a proxy for the measurement of protein activity). A dashed black line shows the "0" activity mark to serve as a reference for catalytically inactive proteins. Panel B shows the mean ± SEM Log $IC_{50}$ (a direct measurement of CTS resistance). Raw data for the three biological replicates of each protein from this are shown in open circles and jittered with respect to the x axis. Raw data from six biological replicates of three proteins (*Leptodactylus*) from Mohammadi et al. [1] are included. Three substitutions produced catalytically inactive proteins, resulting in no measurable $IC_{50}$, and are thus marked by "x" in panel B. A dashed black line shows pig NKA Log$IC_{50}$ and serves as a reference for CTS-sensitivity.

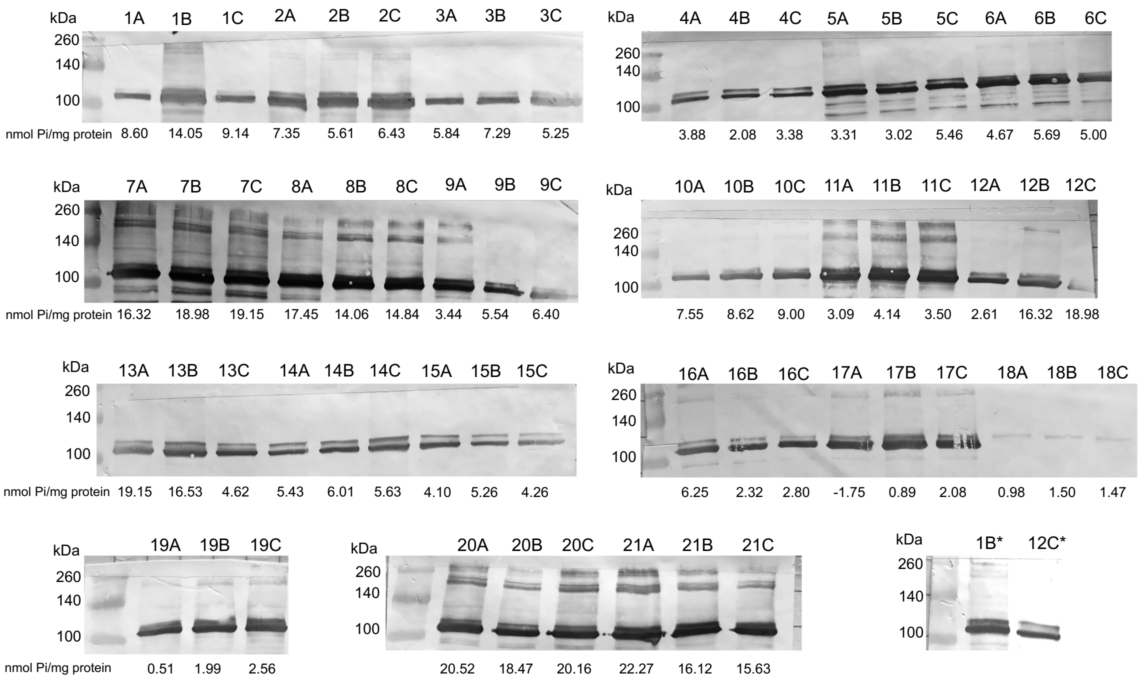


1 – Rat
2 – Chinchilla
3 – False fer-de-lance
4 – Red-necked keelback
5 – Savannah monitor
6 – Black tegu
7 – Ostrich
8 – Yellow-throated sandgrouse
9 – Rat +R111E
10 – Rat +D122H
11 – Chinchilla +N122D
12 – False fer-de-lance +H122D
13 – False fer-de-lance +H122T
14 – False fer-de-lance +H111E
15 – Red-necked keelback +E111H
16 – Savannah monitor +T111H
17 – Savannah monitor +N122H
18 – Black tegu +Q111T
19 – Black tegu +H122D
20 – Ostrich +Q111R
21 – Yellow-throated sandgrouse +R111Q

**Fig. S8.** Western blot analysis of Na,K-ATPase with engineered ATP1A1 (a) subunits produced in this study. The 110 kDa ATP1A1 protein is stained with the α5 monoclonal antibody followed by a horseradish peroxidase conjugated goat antimouse antibody. Samples represents three biological replicates of 21 different recombinant $Na^+$, $K^+$-ATPase (Table S3) produced through cell culture. Protein activity levels (nmol Pi/mg protein) of each biological replicate are indicated under their respective band. Two recombinant proteins (1B* and 12C*) were run a second time on a separate gel due to either the original being cut off on the membrane or the sample concentration was too high.

**Table S1.** Species collection information for data generated by this study. See Supplementary dataset 1 for a complete species list used in this study. Animal purchases were approved by IACUC Protocol No. 2057-16.

| Species | Common name | Source |
|---|---|---|
| *Dendrobates auratus* | Green black poison dart frog | purchased from pet vender |
| *Duttaphrynus melanostictus* | Asian spiny toad | purchased from pet vender |
| *Melanophryniscus stelzneri* | Bumble bee toad | purchased from pet vender |
| *Rhinella granulosa* | Common lesser toad | collected in Colombia |
| *Rhinella marina* | Cane toad | collected in Colombia |
| *Atelopus zeteki* | Panamanian golden frog | collected in Colombia |
| *Atelopus spp* | unknown | collected in Colombia |
| *Phyllomedusa hypochondrialis* | Tiger leg tree frog | purchased from pet vender |
| *Ceratophrys cranwelli* | Pacman frog | purchased from pet vender |
| *Ceratophrys ornatus* | Ornate Pacman frog | purchased from pet vender |
| *Ceratophrys calcarata* | Venezuelan horned frog | collected in Colombia |
| *Lepidobatrachus laevis* | Budgetts frog | purchased from pet vender |
| *Pyxicephalus adspersus* | Pixie frog | purchased from pet vender |
| *Kaloula pulchra* | Chubby frog | purchased from pet vender |
| *Rana catesbeiana* | American bullfrog | purchased from pet vender |
| *Rana sphenocephala* | Leopard frog | purchased from pet vender |
| *Megophrys nasuta* | Long-nosed horned frog | purchased from pet vender |
| *Varanus dumerilii* | Dumerils monitor | purchased from pet vender |
| *Varanus exanthematicus* | Savannah monitor | purchased from pet vender |
| *Varanus niloticus* | Nile monitor | purchased from pet vender |
| *Varanus similis* | Similis monitor | purchased from pet vender |
| *Varanus acanthus* | Spiny-tailed monitor 'Ackie' | purchased from pet vender |
| *Tupinambis teguixin* | Black tegu | collected in Colombia |
| *Ameiva ameiva* | Green ameiva | purchased from pet vender |
| *Rhabdophis subminiatus* | Red-necked keelback snake | obtained from UCB MVZ:Herp: 258317 |
| *Thamnophis sauritus* | Ribbon snake | purchased from pet vender |
| *Heterodon nasicus* | Western hognose snake | purchased from pet vender |
| *Atractus crssicaudtus* | Thickhead ground snake | collected in Colombia |
| *Leptodeira annulata* | Banded cat-eyed snake | collected in Colombia |
| *Leptodeira septentrionalis* | Northern cat-eyed snake | collected in Colombia |
| *Xenodon rhabdocephalus* | False fer-de-lance | collected in Colombia |
| *Helicops angulatus* | Brown-banded water snake | collected in Colombia |
| *Helicops pastazae* | Shreve's keelback | collected in Colombia |
| *Dipsas catesbyi* | Catesby's snail-eater | collected in Colombia |
| *Dendrophidion bivittatus* | Forest racer | collected in Colombia |
| *Oxyrhopus petolarius* | Forest flame snake | collected in Colombia |
| *Diadophis punctatus* | Ring-necked snake | obtained from Dr. Jamie Oaks |
| *Bothrops atrox* | Fer-de-lance | collected in Colombia |
| *Boa constrictor* | Red-tailed boa | collected in Colombia |

**Table S2.** New reptile RNA-seq data generated by this study (PRJNA754197) and amphibian RNA-seq data mined from previous work (PRJNA627222). Refer to Supplementary dataset 1 for sources of ATP1A sequences included in the phylogenetic analysis (Fig. 2)

| Species | Brain | Muscle | Stomach | Skin |
|---|---|---|---|---|
| *Dendrobates auratus* | X | X | X | |
| *Duttaphrynus melanostictus* | X | X | X | |
| *Melanophryniscus stelzneri* | X | X | X | |
| *Rhinella granulosa* | | | X | X |
| *Rhinella marina* | X | | | X |
| *Atelopus zeteki* | | | | X |
| *Atelopus spp* | | | | X |
| *Phyllomedusa hypochondrialis* | X | X | X | |
| *Ceratophrys cranwelli* | X | X | X | |
| *Ceratophrys ornatus* | X | X | X | |
| *Ceratophrys calcarata* | X | X | X | |
| *Lepidobatrachus laevis* | X | X | X | |
| *Pyxicephalus adspersus* | X | X | X | |
| *Kaloula pulchra* | X | X | X | |
| *Rana catesbeiana* | X | X | X | |
| *Rana sphenocephala* | X | X | X | |
| *Megophrys nasuta* | X | X | X | |
| *Varanus dumerilii* | X | X | X | |
| *Varanus exanthematicus* | X | X | X | |
| *Varanus niloticus* | X | X | X | |
| *Varanus similis* | X | X | X | |
| *Varanus acanthus* | X | X | X | |
| *Heloderma suspectum* | X | X | X | |
| *Anolis carolinensis* | X | X | X | |
| *Rhabdophis subminiatus* | X | | | |
| *Nerodia* sp. | X | X | X | |
| *Thamnophis sauritus* | X | X | X | |
| *Thamnophis* sp. | X | X | X | |
| *Heterodon nasicus* | X | X | X | |
| *Atractus crssicaudtus* | | X | X | |
| *Leptodeira annulata* | X | X | X | |
| *Leptodeira septentrionalis* | X | | X | |
| *Xenodon angustirostris* | X | X | X | |
| *Helicops angulatus* | X | X | X | |
| *Helicops pastazae* | X | X | X | |
| *Dipsas catesbyi* | X | X | X | |
| *Dendrophidion bivittatus* | X | | X | |
| *Oxyrhopus petolarius* | X | X | X | |
| *Diadophis punctatus* | X | X | X | |
| *Bothrops atrox* | X | X | | |
| *Boa constrictor* | X | X | X | |

**Table S3.** List of gene constructs used to test functional effects of amino acid substitutions at positions 111 and 122 in vertebrate ATP1A1 (see also Fig S3). For each recombinant protein, the wildtype ATP1B1 of the corresponding species was co-expressed with ATP1A1. Following convention, amino acid positions are based on the sheep numbering system. Wildtype amino acid states at 111 and 122 of ATP1A1 are indicated in parentheses for each species under "Description". Dietary data based on Mohammadi et al. [2]. Data from grass frog (*Leptodactylus macrosternum*) constructs were obtained from Mohammadi et al. [1]

| Construct Name | Engineered Substitution | Description | Known to feed on CTS? |
|---|---|---|---|
| Rat | - | *Rattus norvegicus* wildtype ATP1A1 (R111 & D122) | Y |
| Chinchilla | - | *Chinchilla lanigera* wildtype ATP1A1 (E111 & N122) | N |
| False fer-de-lance | - | *Xenodon rabdocephalus* wildtype ATP1A1 (H111 & H122) | Y |
| Red-necked keelback | - | *Rhabdophis tigrinus* wildtype ATP1A1 (E111 & N122) | Y |
| Savannah monitor | - | *Varanus exanthematicus* wildtype ATP1A1 (T111 & N122) | N |
| Tegu | - | *Tupinambis teguixin* wildtype ATP1A1 (Q111 & H122) | N |
| Ostrich | - | *Struthio camelus* wildtype ATP1A1 (Q111 & N122) | N |
| Yellow-throated sandgrouse | - | *Pterocles gutturalis* wildtype ATP1A1 (R111 & N122) | unknown |
| Rat +R111E | R111E | *R. norvegicus* ATP1A1 + R111E | |
| Rat +D122H | D122H | *R. norvegicus* ATP1A1 + D122H | |
| Chinchilla +N122D | N122D | *C. lanigera* ATP1A1 + N122D | |
| False fer-de-lance +H122D | H122D | *X. rabdocephalus* ATP1A1 + H122D | |
| False fer-de-lance +H111E | H111E | *X. rabdocephalus* ATP1A1 + H111E | |
| False fer-de-lance +H111T | H111T | *X. rabdocephalus* ATP1A1 + H111T | |
| Red-necked keelback +E111H | E111H | *R. tigrinus* ATP1A1 + E111H | |
| Savannah monitor +T111H | T111H | *V. exanthematicus* ATP1A1 + T111H | |
| Savannah monitor +N122H | N122H | *V. exanthematicus* ATP1A1 + N122H | |
| Black tegu +H122D | H122D | *T. teguixin* ATP1A1 + H122D | |
| Black tegu +Q111T | Q111T | *T. teguixin* ATP1A1 + Q111T | |
| Ostrich +Q111R | Q111R | *S. camelus* ATP1A1 + Q111R | |
| Yellow-throated sandgrouse +R111Q | R111Q | *P. gutturalis* ATP1A1 + R111Q | |
| Grass frog | - | *Leptodactylus macrosternum* wildtype ATP1A1 (Q111 & N122) | Y |
| Grass frog +Q111R | Q111R | *L. macrosternum* ATP1A1 + Q111R | |
| Grass frog +N122D | N122D | *L. macrosternum* ATP1A1 + N122D | |

**Table S4.** Summary of the ouabain sensitivity and catalytic properties of $Na^+,K^+$-ATPase for each ATP1A1 recombinant protein construct. The values represent the mean and SD ouabain sensitivity (log $IC_{50}$) of protein activity of three biological replicates. ATP1B1 of each recombinant protein construct was co-expressed with ATP1A1. 'X' indicates that there was no measurable $IC_{50}$. Data from grass frog (*Leptodactylus macrosternum*) constructs were obtained from Mohammadi et al. [1].

| Construct Name | Ouabain sensitivity (mol/L) Mean(log $IC_{50}$) ± SD | Protein activity nmol Pi/(mg protein*min) ± SD |
|---|---|---|
| Rat | -4.420 ± 0.286 | 10.597 ± 3.000 |
| Chinchilla | -5.198 ± 0.207 | 6.464 ± 0.871 |
| False fer-de-lance | -4.055 ± 0.192 | 6.130 ± 1.050 |
| Red-necked keelback | -4.103 ± 0.194 | 2.950 ± 0.750 |
| Savannah monitor | -5.280 ± 0.276 | 3.929 ± 1.336 |
| Tegu | -5.274 ± 0.338 | 5.123 ± 0.521 |
| Ostrich | -5.565 ± 0.086 | 18.153 ± 1.586 |
| Yellow-throated sandgrouse | -4.679 ± 0.005 | 15.448 ± 1.776 |
| Rat +R111E | -6.042 ± 0.777 | 5.127 ± 1.521 |
| Rat +D122H | -4.355 ± 0.185 | 8.392 ± 0.752 |
| Chinchilla +N122D | -2.333 ± 0.495 | 3.577 ± 0.526 |
| False fer-de-lance +H122D | -5.596 ± 0.127 | 13.381 ± 7.835 |
| False fer-de-lance +H111E | -5.257 ± 0.300 | 12.638 ± 8.787 |
| False fer-de-lance +H111T | -5.164 ± 0.354 | 5.691 ± 0.296 |
| Red-necked keelback +E111H | -6.821 ± 0.058 | 4.540 ± 0.627 |
| Savannah monitor +T111H | -4.850 ± 0.332 | 3.790 ± 2.145 |
| Savannah monitor +N122H | X | 1.332 ± 0.309 |
| Black tegu +Q111T | X | 1.695 ± 1.070 |
| Black tegu +H122D | X | 0.409 ± 1.959 |
| Ostrich +Q111R | -4.406 ± 0.259 | 19.718 ± 1.091 |
| Yellow-throated sandgrouse +R111Q | -4.698 ± 0.147 | 18.007 ± 3.699 |
| Grass frog | -5.634 ± 0.586 | 16.295 ± 5.008 |
| Grass frog + Q111R | -4.889 ± 0.851 | 9.926 ± 3.708 |
| Grass frog + N122D | -5.060 ± 0.661 | 7.505 ± 3.623 |

**Table S5.** Tests for background-dependence (epistasis) of substitutions with respect to resistance to ouabain inhibition (IC50) and ATPase activity.

| | IC50 | | | Activity | | |
|---|---|---|---|---|---|---|
| Substitution | Background | Effect | Background *Effect | Background | Effect | Background *Effect |
| H111T/T111H | **0.03** | **0.002** | 0.08 | **0.04** | 0.16 | 0.18 |
| Q111R/R111Q | 0.38 | **0.010** | 0.10 | 0.38 | 0.13 | 0.55 |
| H111E/E111H | **1.4E-4** | **2.1E-4** | **1.8E-7** | **9.0E-4** | **0.04** | 0.21 |
| N122D | **6.6E-5** | **1.9E-4** | **0.0012** | **0.0030** | **0.0022** | 0.16 |
| H122D/D122H | 0.16 | 0.12 | **0.03** | 0.97 | 0.49 | 0.49 |

Notes:
H111T/T111H (false fer-de-lance/Savannah monitor)
Q111R/R111Q (grass frog/ostrich/yellow-throated sandgrouse)
H111E/E111H (false fer-de-lance/red-necked keelback)
N122D (grass frog/chinchilla)
H122D/D122H (false fer-de-lance/black tegu/rat)
Bold values indicate significant raw ANOVA p-values.

**Table S6.** Ancestral sequence reconstruction of aM1-M2 region of the ATP1A1, ATP1A2, ATP1A3 proteins (Fig 2 main text).

| | **Node** | **node_tree** | **node support (aLRT)** | **108** | **111** | **112** | **113** | **114** | **115** | **116** | **117** | **118** | **119** | **120** | **121** | **122** |
|---|---|---|---|---|---|---|---|---|---|---|---|---|---|---|---|---|
| **ATP1A1** | AncAmphibian | 1236 | 2.33 | Y | Q | A | A | T | E | E | E | P | Q | N | D | N |
| **ATP1A1** | AncReptile | 1401 | 7.5 | Y | Q | A | A | T | E | E | E | P | N | N | D | N |
| **ATP1A1** | AncMammal | 1280 | 182.34 | Y | Q | A | A | T | E | E | E | P | Q | N | D | N |
| **ATP1A2** | AncAmphibian | 838 | 282.12 | Y | Q | A | A | M | E | D | E | P | A | N | D | N |
| **ATP1A2** | AncReptile | 866* (paraphyletic) | 0 | Y | Q | A | A | M | E | D | E | P | A | N | D | N |
| **ATP1A2** | AncMammal | 895 | 171.48 | Y | Q | A | A | M | E | D | E | P | S | N | D | N |
| **ATP1A3** | AncAmphibian | 1204 | 102.17 | Y | Q | A | G | M | E | D | D | P | A | G | D | N |
| **ATP1A3** | AncReptile | 1163 | 21.31 | Y | Q | A | G | T | E | D | D | P | S | N | D | N |
| **ATP1A3** | AncMammal | 1034 | 43.12 | Y | Q | A | G | T | E | D | D | P | S | G | D | N |

| | | | | **PP** | | | | | | | | | | | | |
|---|---|---|---|---|---|---|---|---|---|---|---|---|---|---|---|---|
| **ATP1A1** | AncAmphibian | 1236 | 2.33 | 1 | 1 | 1 | 1 | 1 | 1 | 1 | 1 | 1 | 0.587 | 1 | 1 | 1 |
| **ATP1A1** | AncReptile | 1401 | 7.5 | 0.998 | 1 | 1 | 1 | 0.967 | 0.998 | 1 | 1 | 1 | 0.996 | 1 | 1 | 1 |
| **ATP1A1** | AncMammal | 1280 | 182.34 | 1 | 1 | 1 | 1 | 1 | 1 | 1 | 1 | 1 | 1 | 1 | 1 | 1 |
| **ATP1A2** | AncAmphibian | 838 | 282.12 | 1 | 1 | 1 | 1 | 1 | 1 | 1 | 1 | 1 | 0.979 | 1 | 1 | 1 |
| **ATP1A2** | AncReptile | 866* (paraphyletic) | 0 | 1 | 1 | 1 | 1 | 1 | 1 | 1 | 1 | 1 | 0.979 | 1 | 1 | 1 |
| **ATP1A2** | AncMammal | 895 | 171.48 | 1 | 1 | 1 | 1 | 1 | 1 | 1 | 1 | 1 | 0.999 | 1 | 1 | 1 |
| **ATP1A3** | AncAmphibian | 1204 | 102.17 | 1 | 1 | 1 | 1 | 0.974 | 1 | 1 | 1 | 1 | 1 | 1 | 1 | 1 |
| **ATP1A3** | AncReptile | 1163 | 21.31 | 1 | 1 | 1 | 1 | 1 | 1 | 1 | 0.956 | 1 | 1 | 0.935 | 1 | 1 |
| **ATP1A3** | AncMammal | 1034 | 43.12 | 1 | 1 | 1 | 1 | 1 | 1 | 1 | 1 | 1 | 1 | 1 | 1 | 1 |

NOTE – Nodes shown in yellow in Fig. 2A in main text. aLRS: approximate Likelihood-ratio statistic; PP: Posterior probability.

**Table S7.** Summary of the functional comparisons amongst constructs (see Fig 5 in main text). Shown are the 11 pairwise comparisons analyzed: 4 comparisons include states at sites 111 and 7 comparisons at site 122. The % Δ effect of the same amino acid state on two backgrounds (Perc_diff) is shown for each comparison, along with the pairwise number of amino acid differences across the entire protein (AA_dist), the percent pairwise sequence divergence (Perc_diff), and the pairwise number of amino acid differences at the 16 sites identified with ANOVA (see Tables S8 and S9).

| SpeciesA | SpeciesB | AAState | AA_dist | Perc_dist | Perc_diff | AA_dist_16 |
|---|---|---|---|---|---|---|
| Ostrich | Leptodactylus | 111R | 77 | 7.40384615 | 46.0647382 | 1 |
| Rat | Xenodon | 111E | 75 | 7.21153846 | 44.4559222 | 16 |
| Xenodon | Tupinambis | 111T | 49 | 7.40384615 | 191.872877 | 15 |
| Rhabdophis | Varanus | 111H | 52 | 5 | 57.425878 | 10 |
| Chinchilla | Xenodon | 122D | 82 | 7.40384615 | 150.829675 | 16 |
| Chinchilla | Tupinambis | 122D | 76 | 7.30769231 | 22.2484489 | 1 |
| Chinchilla | Leptodactylus | 122D | 86 | 7.40384615 | 8.04020061 | 1 |
| Xenodon | Tupinambis | 122D | 49 | 4.71153846 | 173.078124 | 15 |
| Xenodon | Leptodactylus | 122D | 80 | 7.40384615 | 158.869876 | 16 |
| Tupinambis | Leptodactylus | 122D | 65 | 6.25 | 14.2082483 | 1 |
| Rat | Varanus | 122H | 67 | 7.40384615 | 68.3292549 | 2 |

**Table S8.** Model selection summary for ANOVA nested models and Pearson's correlations under two grouping criteria: Up, sites were grouped according to |ρ| > 0.8 resulting in 24 groups; Down, sites were grouped according to |ρ| > 0.99 resulting in 45 groups. ANOVA models increase in complexity in a stepwise fashion by adding one group at a time, adding groups in the order of the largest (group 1) to smallest (group 24 or group 45) marginal variance explained. Number of sites (# sites) is the cumulative number of sites included in the model. P-values of likelihood ratio tests (p_LRT) and AIC statistic were used for model selection. Note that regardless of the grouping criteria used, the best model includes the same 16 sites (model B and model D, respectively). For each ANOVA model, we used Pearson's correlation to estimate the strength of the relationship between pairwise divergence at the sites included in the model (# sites) and % Δ effect of the same amino acid state on two backgrounds. Because our experimental comparisons are not independent, we performed 10,000 permutations of the experimental pairwise % Δ effect across construct comparisons and estimated the P-value of the model as the probability of observing a Pearson's correlation coefficient higher or equal than the observed.

| Grouped as r>0.8; 24 groups of sites | | | | | | | |
|---|---|---|---|---|---|---|---|
| ANOVA | | | | | | Correlation | |
| **Model** | **group** | **r2_per_model** | **p_LRT** | **aic** | **# sites** | **Pearson's r** | **perm.Pval** |
| A | 1 | 0.5995211 | NA | 121.3514 | 14 | 0.80183079 | 0.0033 |
| B | 1-2 | 0.7752917 | 1.93E-02 | 116.99495 | 16 | 0.7879634 | 0.003 |
| C | 1-3 | 0.8007926 | 3.81E-01 | 117.66991 | 22 | 0.64910697 | 0.0185 |
| D | 1-4 | 0.827243 | 3.82E-01 | 118.10284 | 23 | 0.60719372 | 0.0285 |
| E | 1-5 | 0.8339467 | 6.88E-01 | 119.66749 | 29 | 0.36417096 | 0.1287 |
| F | 1-6 | 0.8508927 | 5.59E-01 | 120.48342 | 32 | 0.31381135 | 0.1694 |
| G | 1-7 | 0.90601 | 2.79E-01 | 117.40713 | 43 | 0.04898121 | 0.4317 |
| H | 1-8 | 0.9986534 | 1.06E-16 | 72.69779 | 45 | 0.0520149 | 0.433 |

Max R^2 = 0.9986534

| Grouped as r>0.99; 45 groups of sites | | | | | | | |
|---|---|---|---|---|---|---|---|
| ANOVA | | | | | | Correlation | |
| **Model** | **group** | **r2_per_model** | **p_LRT** | **aic** | **# sites** | **Pearson's r** | **perm.Pval** |
| A | 1 | 0.6626033 | NA | 119.46598 | 6 | 0.81199806 | 0.0036 |
| B | 1-2 | 0.6850202 | 4.80E-01 | 120.70972 | 14 | 0.80183079 | 0.0023 |
| C | 1-3 | 0.7138161 | 4.37E-01 | 121.6551 | 15 | 0.80595708 | 0.0019 |
| D | 1-4 | 0.827243 | 7.00E-02 | 118.10284 | 16 | 0.7879634 | 0.0025 |
| E | 1-5 | 0.8339467 | 6.88E-01 | 119.66749 | 17 | 0.76557677 | 0.0053 |
| F | 1-6 | 0.8339467 | NA | 119.66749 | 22 | 0.64910697 | 0.0217 |
| G | 1-7 | 0.8339467 | NA | 119.66749 | 23 | 0.60719372 | 0.0305 |
| H | 1-8 | 0.8339467 | NA | 119.66749 | 24 | 0.56420809 | 0.0402 |
| I | 1-9 | 0.8619802 | 4.35E-01 | 119.63345 | 27 | 0.4518028 | 0.0817 |
| J | 1-10 | 0.8619802 | NA | 119.63345 | 29 | 0.36417096 | 0.142 |
| K | 1-11 | 0.9396978 | 1.08E-01 | 112.52503 | 30 | 0.32288166 | 0.1621 |
| L | 1-12 | 0.9986534 | 3.61E-11 | 72.69779 | 31 | 0.29173786 | 0.199 |

Max R^2 = 0.9986534

**Table S9.** 16 sites included in the two groups (r>0.8) that explain a high proportion of variation in functional effects identified by ANOVA modeling. F-values, R^2, and P-values correspond to individual ANOVA analysis per site.

| site | Fvals | R2 | pvals | LogP | group |
|---|---|---|---|---|---|
| 226 | 15.7110034 | 0.6626033 | 0.00415537 | 5.48335344 | 1 |
| 271 | 15.7110034 | 0.6626033 | 0.00415537 | 5.48335344 | 1 |
| 290 | 15.7110034 | 0.6626033 | 0.00415537 | 5.48335344 | 1 |
| 406 | 15.7110034 | 0.6626033 | 0.00415537 | 5.48335344 | 1 |
| 645 | 15.7110034 | 0.6626033 | 0.00415537 | 5.48335344 | 1 |
| 832 | 15.7110034 | 0.6626033 | 0.00415537 | 5.48335344 | 1 |
| 102 | 11.9761117 | 0.59952112 | 0.00855958 | 4.76070454 | 1 |
| 179 | 11.9761117 | 0.59952112 | 0.00855958 | 4.76070454 | 1 |
| 224 | 11.9761117 | 0.59952112 | 0.00855958 | 4.76070454 | 1 |
| 227 | 11.9761117 | 0.59952112 | 0.00855958 | 4.76070454 | 1 |
| 416 | 11.9761117 | 0.59952112 | 0.00855958 | 4.76070454 | 1 |
| 521 | 11.9761117 | 0.59952112 | 0.00855958 | 4.76070454 | 1 |
| 888 | 11.9761117 | 0.59952112 | 0.00855958 | 4.76070454 | 1 |
| 979 | 11.9761117 | 0.59952112 | 0.00855958 | 4.76070454 | 1 |
| 561 | 8.89454233 | 0.52647382 | 0.01753683 | 4.04345233 | 2 |
| 279 | 7.74965712 | 0.49205198 | 0.02378394 | 3.73874456 | 2 |
